# OPT3 Transports Copper to the Phloem, Mediates Shoot-to-Root Copper Signaling and Crosstalk Between Copper and Iron Homeostasis in *A. thaliana*

**DOI:** 10.1101/2021.07.30.454504

**Authors:** Ju-Chen Chia, Jiapei Yan, Maryam Rahmati Ishka, Marta Marie Faulkner, Eli Simons, Rong Huang, Louisa Smieska, Arthur Woll, Ryan Tappero, Andrew Kiss, Chen Jiao, Zhangjun Fei, Leon V. Kochian, Elsbeth Walker, Miguel Piñeros, Olena K. Vatamaniuk

## Abstract

Copper and iron are essential micronutrients but are toxic when accumulating in cells in excess. Thus, their uptake by roots is tightly regulated. While plants sense and respond to local copper availability, the systemic regulation of copper uptake has not been documented. By contrast, both local and systemic control for iron uptake has been reported. Iron abundance in the phloem has been suggested to act systemically, regulating the expression of iron uptake genes in the root. Consistently, shoot-to-root iron signaling is disrupted in *A. thaliana* mutants lacking the phloem companion cell-localized iron transporter, AtOPT3: *opt3* mutants overaccumulate iron in leaves while constitutively upregulating iron deficiency-responsive genes in roots. We report that AtOPT3 transports copper and mediates its delivery from source leaves to sinks including young leaves and developing embryos. Consequently, the *opt3* mutant accumulates less copper in the phloem, is sensitive to copper deficiency, and mounts transcriptional copper deficiency response in roots. Copper rescues these defects. Notably, feeding the *opt3* mutant with copper or iron *via* the phloem in leaves downregulates the expression of both copper and iron-deficiency marker genes in roots, suggesting that copper and iron can substitute each other’s function in the phloem in shoot-to-root communication.

**One-sentence summary:** AtOPT3 loads copper and iron into the phloem companion cells, for subsequent distribution to sink tissues and systemic signaling of copper and iron deficiency.

## Introduction

Iron and copper are essential elements, required in trace amounts to complete the life cycle of all organisms, including plants and humans. However, these elements are toxic to cells if they accumulate in ionic form (Broadley et al., 2012; Ravet and Pilon, 2013). The essential yet toxic nature of iron and copper is attributed to the ease with which they accept and donate electrons (Broadley et al., 2012; Ravet and Pilon, 2013). This ability has been capitalized by nature, and so copper and iron-containing enzymes are required for vital physiological processes, including photosynthesis, respiration and scavenging of reactive oxygen species (Broadley et al., 2012; Ravet and Pilon, 2013). In addition to these processes, copper is required for cell wall lignification and reproduction (Epstein and Bloom, 2005; Broadley et al., 2012; Chen et al., 2020; Rahmati Ishka and Vatamaniuk, 2020; Sheng et al., 2021; Rahmati Ishka et al., 2022). Recent studies implicate copper in light-dependent seed germination (Jiang et al., 2020), shaping the shoot architecture, transition to flowering, stigmatic papillae development and senescence (Rahmati Ishka and Vatamaniuk, 2020; Sheng et al., 2021) and reviewed in (Rahmati Ishka et al., 2022). Mounting evidence from studies in animal species suggests that in addition to a static function as a cofactor of cellular enzymes, copper is involved in cell signaling (Turski and Thiele, 2009; Turski et al., 2012; Chang, 2015; Tsang et al., 2020). In plants, copper participates in hormone signaling and accumulation. Specifically, Cu(I) presence in the transmembrane sensor domain of the ethylene receptor, ETR1, is essential for ethylene binding and receptor function in ethylene signaling in *A. thaliana* (Rodríguez et al., 1999; Schott-Verdugo et al., 2019). Likewise, the binding of the plant defense hormone salicylic acid to its receptor NPR1 occurs *via* copper and copper deficiency increases the accumulation of jasmonic acid in leaves of *A. thaliana* (Wu et al., 2012; Yan et al., 2017; Rahmati Ishka et al., 2022). Iron-containing enzymes are also involved in nitrate and sulfate assimilation and chlorophyll synthesis, as well as ethylene and jasmonic acid accumulation (Broadley et al., 2012; Li and Lan, 2017; Cui et al., 2018).

Copper and iron uptake by plant roots and their internal transport and storage are rigorously regulated at transcriptional and posttranscriptional levels in response to the availability of these elements in a local environment and the demands of the developing shoot (Burkhead et al., 2009; Ravet and Pilon, 2013; Kobayashi, 2019; Pottier et al., 2019; Spielmann and Vert, 2020). To maintain copper homeostasis, plants regulate cellular copper uptake and economize on copper by prioritizing it during deficiency from non-essential to essential copper enzymes (Rahmati Ishka et al., 2022). In *A. thaliana*, both processes are controlled by a conserved transcription factor, SPL7 (**S**quamosa **P**romoter Binding Protein–**l**ike7 (Yamasaki et al., 2009; Bernal et al., 2012). In addition to SPL7, a member of the basic helix-loop-helix (bHLH) family, CITF1 (**C**opper Deficiency-**i**nduced **T**ranscription **F**actor 1, *alias* bHLH160) regulates copper uptake into roots, delivery to leaves, flowers and anthers (Yan et al., 2017). CITF1 acts together with SPL7, and the function of both is required for copper delivery to reproductive organs and fertility (Yan et al., 2017). The SPL7-dependent regulon includes Iron (**F**e)/Cu **r**eductase **o**xidases, *FRO4* and *FRO5*, and several copper transporters, including *COPT1, COPT2* and, in part, *COPT6*, that are members of the CTR/COPT/SLC31 (**C**opper **T**ransporter/**Cop**per **T**ransporter/**S**o**l**ute **C**arrier **31**) family (Yamasaki et al., 2009; Bernal et al., 2012; Jung et al., 2012; Gayomba et al., 2013; Jain et al., 2014; Araki et al., 2018; Alexander et al., 2019). *FRO4/5* and *COPT2* are also downstream targets of CITF1 (Yan et al., 2017). Altered expression of SPL7- and CITF1-regulated genes and the increased expression of *CITF1* constitute a signature of the copper deficiency response. Recent studies have shown that SPL7 is expressed mainly in the vasculature of *A. thaliana* and locally regulates root and shoot responses to copper deficiency (Araki et al., 2018). The existence of long-distance copper signaling has not been yet reported.

Regulation of iron homeostasis in *A. thaliana* involves a network of transcription factors from the bHLH family (Jeong et al., 2017; Kim et al., 2019). Specifically, a member of the IVb subgroup of the bHLH family, URI (Upstream Regulator of IRT1, bHLH121), acts upstream as an iron-dependent switch (Kim et al., 2019; Gao et al., 2020). It heterodimerizes with a subgroup IVc bHLH members to regulate the expression of the master regulator of iron homeostasis FIT (**F**ER-like iron deficiency-induced **t**ranscription factor, alias bHLH29) (Kim et al., 2019; Gao et al., 2020). FIT forms heterodimers with the Ib subgroup of bHLH transcription factors to regulate the expression of multiple genes in *A. thaliana* roots, among which are some components of the iron uptake system, *IRT1* (**I**ron-**R**egulated **T**ransporter **1**), *FRO2* and *AHA2* (Plasma Membrane Proton ATPase) (reviewed in (Jeong et al., 2017; Schwarz and Bauer, 2020). The upregulated expression of *Ib bHLH* genes, *FIT*, *AHA2*, *FRO2* and *IRT1*, and the newly discovered *IMA/FEP* iron-sensing peptides (**I**ron **Ma**n [IMA]/**Fe** Uptake-Inducing **P**eptide [FEP]) is a hallmark of root iron-deficiency response (Jeong et al., 2017; Grillet et al., 2018; Hirayama et al., 2018; Schwarz and Bauer, 2020). These and other iron-responsive genes are upregulated in response to local and systemic iron status signals (Gayomba et al., 2015; Jeong et al., 2017; Grillet et al., 2018; Hirayama et al., 2018; Schwarz and Bauer, 2020).

Several mutants with disrupted shoot-to-root iron deficiency signaling have been identified in *A. thaliana* and other plant species, and all exhibit constitutive activation of iron-acquisition genes even when grown under iron-sufficient conditions (Kumar et al., 2017) and reviewed in (Gayomba et al., 2015). Of these, a member of the OPT (**O**ligo **P**eptide **T**ransporter) clade of the OPT transporter family, *A. thaliana* OPT3, is considered a key player in the systemic signaling of iron deficiency (Stacey et al., 2002; Stacey et al., 2008; Mendoza-Cózatl et al., 2014; Zhai et al., 2014). While the *opt3-1* knockout allele is embryo-lethal, the *opt3-2* and *opt3-3* alleles that possess residual levels of *OPT3* expression, overaccumulate iron in roots and leaves because they cannot appropriately downregulate the expression of *AHA2*, *IRT1* and *FRO2* and other iron deficiency-responsive genes (Stacey et al., 2002; Stacey et al., 2008; Mendoza-Cózatl et al., 2014; Zhai et al., 2014). AtOPT3 mediates iron ions uptake into *Xenopus laevis* oocytes and *Saccharomyces cerevisiae*, localizes to companion cells of the phloem, mediates iron loading to the phloem, and facilitates iron delivery from sources (mature leaves) to sinks (roots and seeds) (Zhai et al., 2014). Based on these findings it is suggested that by loading iron into the phloem in leaves, OPT3 communicates iron sufficiency status to the root. Consistent with this suggestion, the loss of the iron transport function in the *opt3-3* mutant and the decreased iron accumulation in its phloem sap not only leads to a decreased iron accumulation in seeds but also is perceived as the iron deficiency signal by the root (Zhai et al., 2014). Local sensing of high iron status in the shoot seems not to be disrupted in the *opt3* mutant, as evidenced by the increased expression of genes encoding iron-storage proteins *FER3* and *FER4* (Khan et al., 2018).

Iron abundance or a lack thereof in plant tissues is tightly linked to the accumulation of other transition metals (Baxter et al., 2008). In this regard, crosstalk between iron and copper is now well documented. The hallmark of this crosstalk is the overaccumulation of copper under iron deficiency and overaccumulation of iron under copper deficiency (Bernal et al., 2012; Waters and Armbrust, 2013; Kastoori Ramamurthy et al., 2018; Rai et al., 2021; Sheng et al., 2021). Consistent with the increased copper uptake under iron deficiency, iron deficiency downregulates the expression of copper-deficiency-regulated genes (Waters et al., 2012; Waters et al., 2014; Yan et al., 2017). On the other hand, iron overaccumulation can also reduce copper uptake in *A. thaliana* and animal systems (Klevay, 2001; Waters and Armbrust, 2013; Ha et al., 2016). Together, these studies suggest that copper and iron homeostasis are interconnected and deficiency for one of the metals or both can interfere with their local and/or long-distance status signaling

Here we provide evidence that in addition to iron, AtOPT3 mediates copper uptake into *Xenopus* oocytes and *S. cerevisiae* cells. Loss of this function in the *opt3-3* mutant results in a decreased copper accumulation in the phloem and reduced copper recirculation from sources (mature leaves) to sinks (roots, young leaves, siliques and developing embryos) compared to wild type. In addition, the *opt3-3* mutant experiences copper deficiency as evidenced by low copper accumulation in roots and young leaves, and increased expression of copper deficiency marker genes. These defects are rescued by copper application. Furthermore, copper feeding *via* the phloem in the shoot rescued molecular symptoms of copper deficiency in the root of wild type and the mutant, highlighting the existence of long-distance shoot-to-root copper signaling. Interestingly, phloem feeding with copper in the shoot also rescued molecular symptoms of iron deficiency in the root of the *opt3-3* mutant and decreased the transcript abundance of molecular markers of iron deficiency in the root of wild type. Likewise, phloem feeding with iron in the shoot downregulated the expression of both iron and copper-deficiency marker genes in the root of the *opt3*-*3* mutant. These data assign new transport capabilities to AtOPT3, provide evidence for the existence of shoot-to-root signaling of copper status, and increase understanding of the crosstalk between copper and iron in long-distance signaling.

## Results

### The *opt3-3* Mutant Accumulates Less Copper in the Phloem

AtOPT3 localizes to the phloem (**Figure 1A** and Zhai et al., 2014), resides in companion cells (Mustroph et al., 2009), and facilitates iron accumulation in the phloem perhaps *via* xylem to phloem transfer (Zhai et al., 2014). As a result of this function, iron concentration in the phloem sap is significantly lower, while it is significantly higher in the xylem sap in the *opt3-3* mutant compared to wild type (Zhai et al., 2014). This finding suggested that the distribution of iron in the vascular tissue of the mutant *vs.* wild type might be altered. To test this hypothesis, we used synchrotron x-ray fluorescence (XRF) microscopy to compare the spatial distribution of iron and other elements in the vasculature of the *opt3-3* mutant (from here on referred to as *opt3*) *vs.* wild type. We first evaluated mineral distribution in mature leaves that serve as photosynthetic sources of nutrients for developing leaves at the vegetative stage. Consistent with our past findings (Zhai et al., 2014), the *opt3* mutant accumulated more iron throughout the leaf blade, with the bulk of iron located in minor veins compared to wild type (**Figure 1B** and **Supplemental Figure 1 online A to D**). We also found that in addition to iron, mature leaves of the *opt3* mutant accumulated more copper (**Figure 1C** and **Supplemental Figure 1 online A to D**), manganese and zinc (**Figure 1C** to **E**). The spatial distribution of manganese and zinc was not altered in the *opt3* mutant compared to wild type: manganese was spread throughout the leaf blade with the highest accumulation in basal cells of trichomes, while zinc was also noticeable in the vasculature (**Figure 1D, E and K**). Copper also overaccumulated in the vasculature of the *opt3* mutant, and its distribution pattern in minor veins resembled the distribution of iron (**Figure 1B, C and K,** and **Supplemental Figure 1 online A to D**).

**Figure 1.**
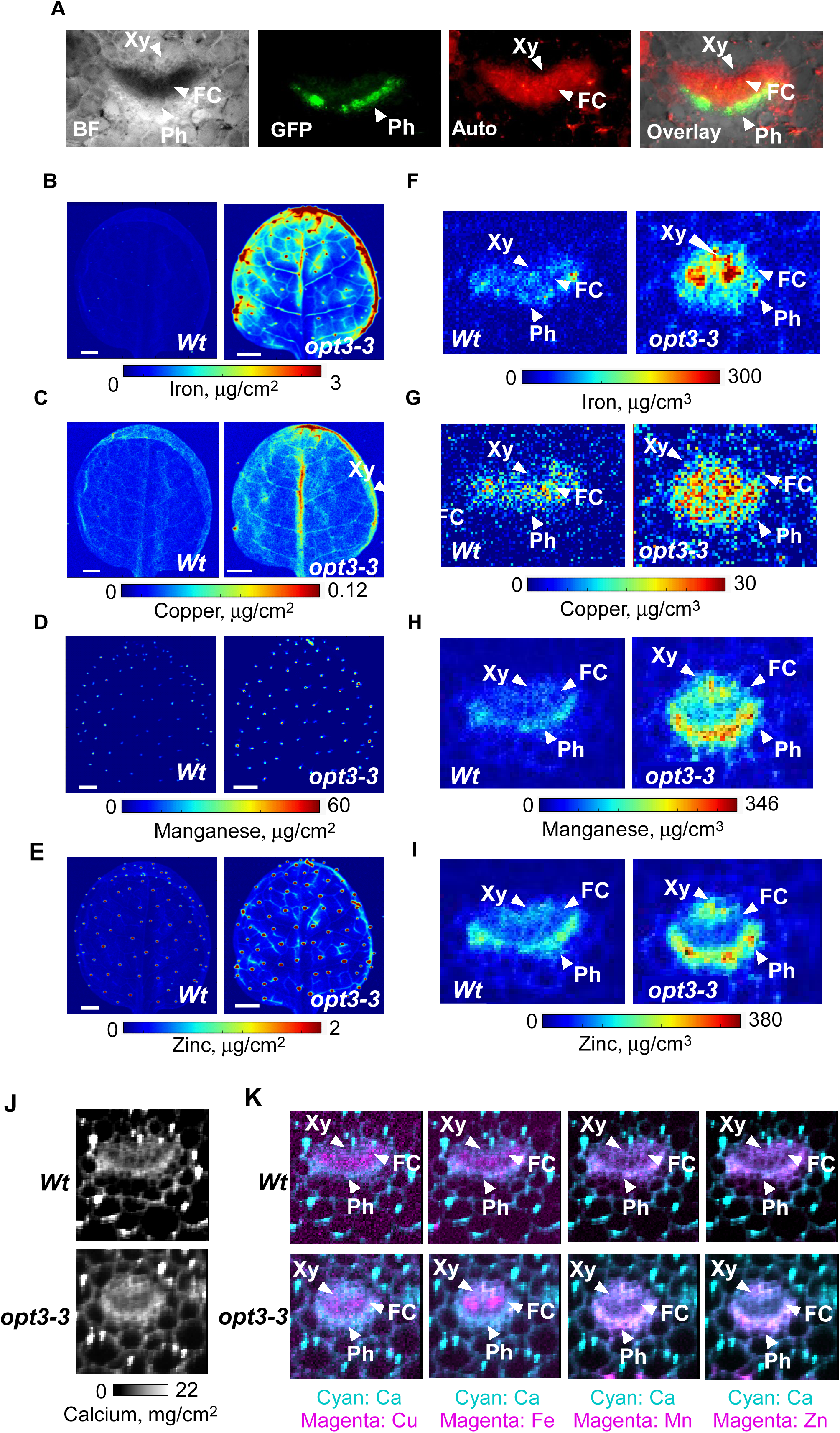
OPT3 mediates copper loading to the phloem. (A) Hand-cut cross sections through the petiole of transgenic wild-type plants expressing the OPT3pro-GFP construct (Zhai et al. 2014). **B** to **E** show 2D XRF maps of the indicated minerals in mature leaves. **F** to **J** show 2D-CXRF maps of indicated elements in the vasculature of mature leaf petioles. (**K**) Merged images of calcium (Ca) and indicated elements. Calcium maps show the overall structures of the vascular tissues, and the majority of copper and iron in the *opt3* mutant is associated with the xylem and fascicular cambium (**FC**). Plants were grown hydroponically with 250 nM CuSO_4_ for 26 days (in **B** to **E**) or five weeks (in **F** to **I**) before tissues were collected In **B** to **K** bar = 1 mm. Xy, FC and Ph indicate the location of the xylem, fascicular cambium and the phloem, respectively. **B** to **K** show representative images from three analyzed plants. Results from two other independent experiments are shown in **Supplemental Figure 1C, D online**).

We then used 2D-XRF in a confocal mode (2D-CXRF) with a specialized x-ray collection optic to obtain a micron-scale resolution; this enabled analyses of mineral localization in the phloem *vs.* xylem in the *opt3* mutant *vs.* wild type. For the current study, this technique is preferable to traditional XRF methods (both 2D XRF and 3D micro-XRF tomography) because it allows quantitative comparisons of metal distributions among different samples without the need to control or limit the sample thickness or lateral size (Mantouvalou et al., 2012). Using 2D-CXRF, we found that the spatial distribution of iron and copper but not of manganese or zinc was altered in the *opt3* mutant compared to wild type (**Figure 1 F** to **I**). This was further confirmed by comparing the distribution of fluorescence from calcium (used to visualize cell boundaries, [**Figure 1J**]) overlayed with the fluorescence from iron or copper, zinc or manganese (**Figure 1K**). Specifically, while iron was evenly distributed between the xylem and the phloem regions in the wild type, the bulk of iron was found in the xylem and fascicular cambium region and to a lesser extent in the phloem region in the *opt3* mutant. (**Figure 1F and K**). Copper was mainly located in the fascicular cambium and phloem regions and, to a lesser extent, the xylem in the wild type (**Figure 1G and K**). By contrast, the bulk of copper was associated with the xylem and fascicular cambium and to a lesser extent, the phloem tissues in the *opt3* mutant (**Figure 1G and K**). The loss of OPT3 function led to a significant increase in the concentration of manganese and zinc in both the xylem and the phloem of the mutant compared to wild type (**Figure 1H, I and K**).

Consistent with 2D-CXRF results, the concentration of copper in the phloem was significantly lower in the *opt3* mutant compared to the wild type (**Figure 2A**). The decreased accumulation of copper in the phloem of the *opt3* mutant *vs.* wild type was independently found by the Walker lab (**Supplemental Fig. S1E online**).

**Figure 2.**
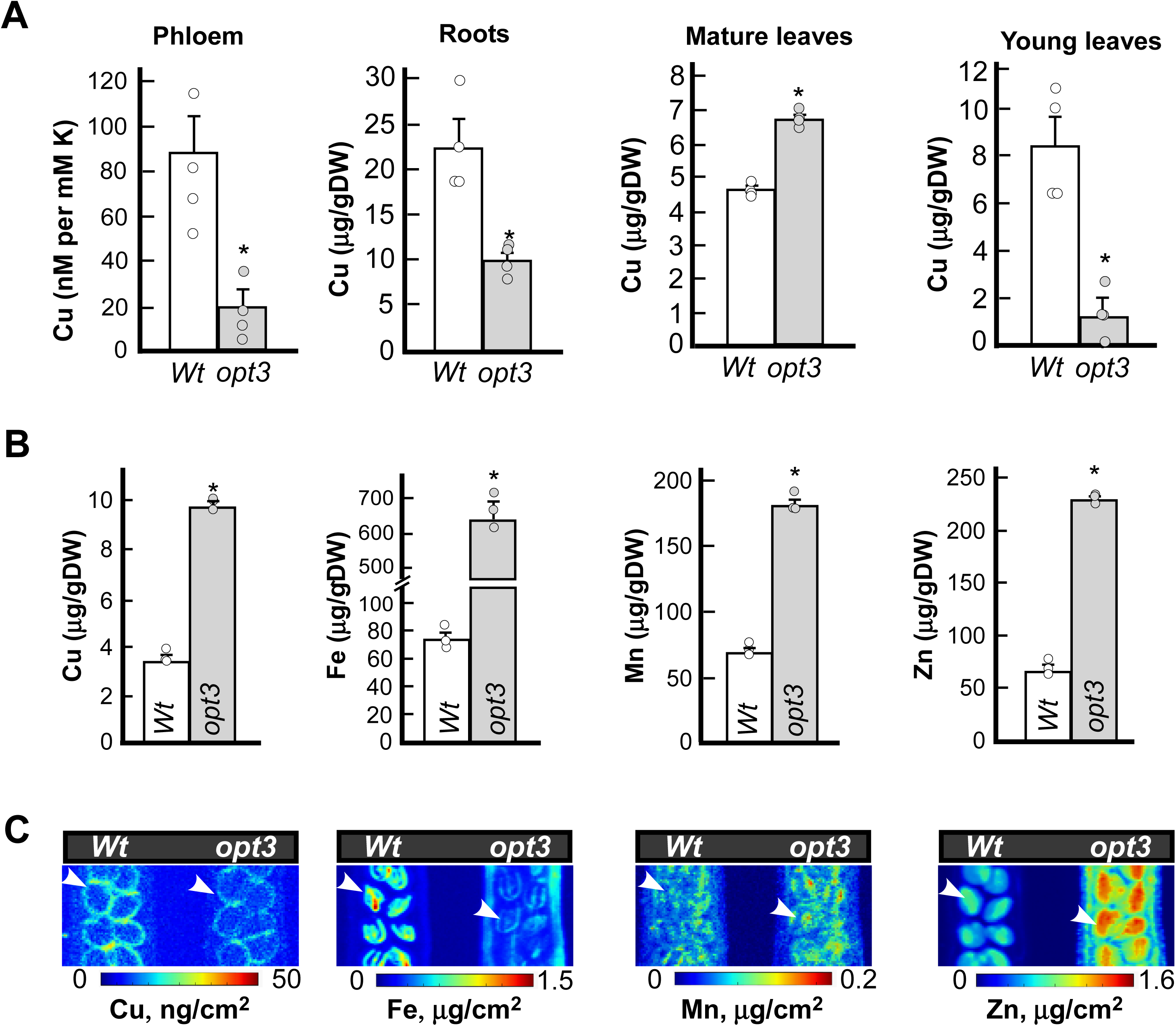
The *opt3* mutant accumulates less copper in the phloem sap, roots, young leaves and developing embryos. (**A**) The concentration of copper in the phloem sap and the indicated plant tissues of wild type and the *opt3* mutant. Plants were grown hydroponically under iron and copper sufficient conditions. Shown values are arithmetic means ± S.E. Asterisks indicate statistically significant differences from wild type (*p* < 0.05, Student’s *t* test, n = 4 independent phloem sap collections). An independent analysis of copper concentration in the phloem sap is shown in **Supplemental Figure 1E online**. (**B**) ICP-MS analysis of mineral accumulation in dry silique valves collected from soil-grown plants. Shown values are arithmetic means ± S.E. Asterisks indicate statistically significant differences from wild type (p < 0.05, Student’s t test, n = 3 independent experimental set-ups. In each experimental set-up, tissues from four to five plants grown in the same container were pooled and represented an independent measurement. Data for the wild type *vs.* the *opt3-2* allele are shown in **Supplemental Figure 3 online**. (**C**) Fifteen-mm-long developing siliques were collected from soil-grown plants and subjected to 2D-SXRF analysis. White arrows point to embryos. A representative image from three independent experiments is shown; results from another independent experiment are shown in **Supplemental Figure 4A online**.

### Roots, Young Leaves and Developing Embryos of the *opt3* Mutant Accumulate Less Copper

Past studies have shown that mutant alleles of *OPT3* over accumulate iron, manganese and zinc in roots and leaves (Stacey et al., 2008; Mendoza-Cózatl et al., 2014; Zhai et al., 2014). Since the 2D-CXRF analysis of mineral distribution in the vasculature has also pointed to the role of OPT3 in copper homeostasis, we refined our past analysis of total internal metal accumulation to include copper. Consistent with past findings, roots and both mature and young leaves of the *opt3* mutant accumulated significantly more iron, manganese and zinc compared to the corresponding organs of wild type (**Supplemental Fig. S2 online**). By contrast, the concentration of copper in the roots of the *opt3* mutant was reduced to less than 1/3 of the wild type level (**Figure 2A**). We also found that the copper concentration was higher in mature leaves (sources) and lower in young leaves (sinks) in the *opt3* mutant *vs.* wild type (**Figure 2A**). Similar results were obtained using a different, *opt3-2* allele (**Supplemental Fig. S3 online**).

We then analyzed copper and also the accumulation and distribution of iron, manganese and zinc in other source tissues such as silique valves and their sinks, developing embryos and seeds. ICP-MS analysis disclosed that silique valves of the *opt3* mutant overaccumulated copper, iron, manganese and zinc compared to wild type (**Figure 2B**). Using 2D-XRF we found that developing embryos of the mutant accumulated less copper and iron (**Figure 2C and Supplemental Figure 4A online**) but more manganese and zinc (**Figure 2C**), suggesting that copper and iron delivery from sources to sinks is reduced in the mutant *vs.* wild type. It was noticeable that the 2D-XRF detectable spatial distribution of copper in developing embryos of both wild type and the *opt3* mutant was distinct from iron. Specifically, copper was associated primarily with the developing seed coat, while iron was mostly localized in the embryo vasculature.

We then used high-resolution 2D computed tomography XRF (CT-XRF) to visualize minerals in mature seeds. Similar to embryos, copper was associated mainly with the seed coat and was detected throughout the seed and the vasculature (**Supplemental Figure 4B online)**. In some areas of the seed coat, copper concentration was lower in the wild type than in the mutant, while the level of copper in the vasculature was lower in the mutant than in the same areas of wild type (**Supplemental Figure 4B online)**. This subtle difference in copper distribution in mutant *vs.* wild type seeds translated to the overall similar internal seed copper concentration in both genotypes (**Supplemental Figure 4C online)**. As was shown previously, iron was associated with the vascular parenchyma cells in mature wild type seeds (**Supplemental Figure 4B online** and (Kim et al., 2006)). Iron distribution did not change in the mutant, although iron accumulation in the *opt3* vascular parenchyma cells was significantly lower than in wild type. Consistently, the total internal iron concentration was lower in the *opt3* mutant seeds *vs.* wild type (**Supplemental Figure 4C online)**. While the distribution of manganese and zinc did not change in the mutant *vs.* wild type, the *opt3* mutant seeds accumulated significantly more zinc (**Supplemental Figure 4B, C online)**. Together, our data suggest that OPT3 contributes to the phloem-based redistribution of copper from mature leaves to young leaves and from silique valves to developing embryos, and thus, in addition to iron, may also transport copper.

### OPT3 Mediates Copper Uptake in *Xenopus* oocytes and *S. cerevisiae*

The ability of OPT3 to transport copper was studied in *Xenopus laevis* oocytes as we have previously shown that in this expression system, OPT3 localizes to the plasma membrane and mediated iron and cadmium ions uptake into oocytes (Zhai et al., 2014). As potential transport substrates, we tested Cu^2+^ (provided as CuSO_4_) and copper complexed with its biological ligand, nicotianamine (Cu-NA). We found that OPT3 transported both free copper ions as well as copper provided as Cu-NA complex into oocytes (**Figure 3A**). Furthermore, OPT3-expressing oocytes accumulated 4.2 times more copper when it was provided as a free ion than when complexed with NA. This finding suggested that free copper ions are a preferred OPT3 substrate, at least in this heterologous system. Visual examination and membrane potential measurements allowed us to assess the cellular integrity of mock and *OPT3* expressing oocytes throughout the uptake experiments to rule out the possibility that the observed fluxes were the product of membrane leakiness due to cellular toxicity rather than specific OPT3-mediated uptake (**Supplemental Figure 5 online)**. Under control conditions, the cells expressing OPT3 had less negative membrane potential than mock cells, consistent with our earlier reports (Zhai et al., 2014). The appearance and survival rates of mock and OPT3-expressing oocytes remained the same within the three hours of the experiment, and only a minor reduction in resting potential (7 and 14 mV in mock and OPT3-expressing cells, respectively) was recorded, suggesting the cells-maintained membrane integrity throughout the uptake time (**Supplemental Figure 5A online)**. After transferring cells into the copper-free solution, the survival rates and resting potentials of OPT3-expressing cells, but not of mock-treated cells, were dramatically reduced over the next 21 h of incubation in the copper-free medium (**Supplemental Figure 5B online**). We concluded that the drop in the survival rates of OPT3-expressing cells resulted from the OPT3-mediated copper-accumulation, leading to cellular toxicity, which was manifested as a loss in the cell’s ability to maintain their resting potential, and eventual death.

**Figure 3.**
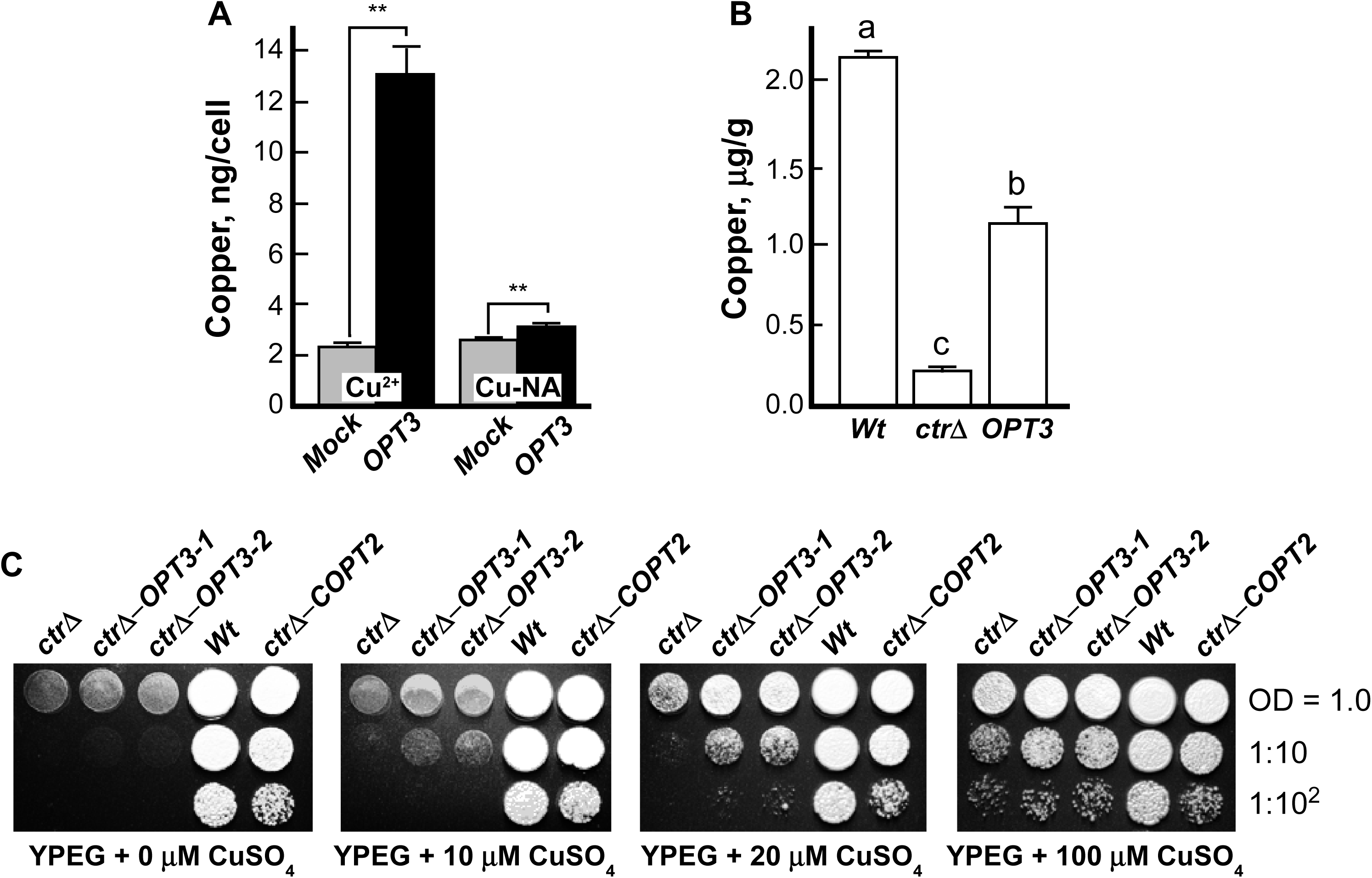
OPT3 transports copper in *X. laevis* oocytes and *S. cerevisiae*. (**A**) Copper uptake into X*enopus* oocytes injected with either *OPT3* cRNA (**OPT3**) or water (**Moc**k). Copper uptake was measured at 3 h. The uptake solution was supplemented with 25 µM Cu-NA (**Cu-NA**) or 100 µM CuSO_4_ (**Cu^2^**^+^, the final free Cu^2+^ activity in the uptake solution being estimated to be 35 µM as determined by GEOCHEM-EZ (Shaff et al., 2010)). Presented values are arithmetic means ± S.E. (n = 5 independent measurements). Asterisks indicate statistically significant differences (**, *p* < 0.01, using Student’s *t*-test). **(B)** Copper concentration in *S. cerevisiae* wild-type, SEY6210 and its isogenic *ctr1Δ2Δ3Δ* mutant both expressing the empty *YES3-Gate* vector (***Wt*** and ***ctrΔ***, respectively) or the *ctr1Δ2Δ3Δ*mutant expressing *YES3-Gate-OPT3* (***ctrΔ−*OPT3**) all grown with 20 µM CuSO_4_. Levels not connected with the same letter are statistically different (n = 4 to 5 independent ICP-MS measurements [independently grown cells]), Tukey HSD test). (**C**) The wild type and the *ctr1Δ2Δ3Δ S. cerevisiae* mutant transformed with the empty YES3-Gate vector or the *ctr1Δ2Δ3Δ* transformed with the vector containing the *OPT3* cDNA or the *A. thaliana* copper transporter, *COPT2*, were serially 10-fold diluted and spotted onto solid YPEG medium supplemented with different concentrations of CuSO_4_. Colonies were visualized after incubating plates for three days at 30°C. Dilution series are indicated on the right. *S. cerevisiae* lines were designated as in (**B**), except that ***OPT3-1*** and ***OPT3-2*** designate two distinct *OPT3* expressing clones selected and propagated after yeast transformation.

The ability of OPT3 to transport copper was further validated by functional complementation assays in the *S. cerevisiae*, which lack the capability to synthesize NA. We used a copper-deficient *S. cerevisiae ctr1Δctr2Δctr3Δ* mutant lacking the high-affinity plasma membrane copper uptake transporters Ctr1p and Ctr3p and the vacuolar membrane copper efflux transporter, Ctr2p (Dancis et al., 1994; Rees et al., 2004). Due to low internal copper, the *ctr1Δctr2Δctr3Δ* mutant cells manifest a respiratory defect because of the altered activity of the copper-dependent cytochrome *c* oxidase complex of the mitochondrial respiratory chain. This defect can be visualized by the failure of the *ctr1Δctr2Δctr3Δ* mutant to grow on non-fermentable carbon sources such as ethanol and glycerol (YPEG medium) unless copper is supplied exogenously (Dancis et al., 1994). As expected, the *ctr1Δctr2Δctr3Δ* cells expressing the empty YES3-Gate vector accumulated 10-fold less copper than the vector-expressing wild type (**Figure 3B**). The expression of *OPT3* in mutant cells increased their copper accumulation by 5-fold compared to the vector expressing cells, although it did not bring it to the level of the vector-expressing wild type cells (**Figure 3B**).

Growth of *ctr1*Δ*ctr2*Δ*ctr3*Δ mutant and wild type strains expressing the empty *YES3-Gate* vector and the *ctr1*Δ*ctr2*Δ*ctr3*Δ mutant transformed with *YES3-Gate* with the *OPT3* cDNA insert was also compared on a medium containing the non-fermentable carbon sources, ethanol and glycerol (YPEG). The *ctr1*Δ*ctr2*Δ*ctr3*Δ mutant transformed with the *A. thaliana* copper transporter, *COPT2*, was used as an additional positive control (Gayomba et al., 2013). As shown previously, the vector-expressing *ctr1*Δ*ctr2*Δ*ctr3*Δ cells did not grow in YPEG medium even when the medium was supplemented with low (10 and 20 µM) concentrations of copper but grew well on YPEG supplemented with 100 µM CuSO_4_ (**Figure 3C** and (Gayomba et al., 2013)). Unlike *COPT2* expressing cells, *OPT3* expressing mutant cells did not grow on YPEG medium without supplemental copper. However, in contrast to vector expressing *ctr1*Δ*ctr2*Δ*ctr3*Δ cells, *OPT3* expressing cells were able to grow when 10 or 20 µM CuSO_4_ was added to the medium (**Figure 3C)**. These results are consistent with the role of OPT3 in copper uptake and suggest that, unlike CTR/COPTs, OPT3 might be a low-affinity copper transporter at least when expressed in *S. cerevisiae*. Together, our results show that OPT3 mainly contributes to the transport of copper ions in heterologous systems.

### The *opt3* Mutant is Sensitive to Copper Deficiency

We next tested the sensitivity of the *opt3* mutant to copper deficiency by comparing its growth and development to the wild type, both grown hydroponically with or without copper supplementation (**Figure 4)**. As we observed previously, the rosette size of the *opt3* mutant was smaller than that in wild type even under control conditions (**Figure 4A** and (Zhai et al., 2014)) and decreased further under copper deficiency *vs.* control conditions; by contrast, copper deficiency did not affect the rosette size of the wild type (**Figure 4A**) early in the vegetative stage of the development. A different *opt3-2*, allele exhibited similar sensitivity to copper deficiency in the medium (**Supplemental Figure S6A**). As evidenced by the shorter root length and lower fresh weight of the *opt3* mutant, the increased sensitivity of the *opt3-3* mutant *vs.* wild type to copper deficiency was also observed in seedlings grown on a solid medium supplemented with the copper chelator bathocuproine disulfonate (BCS) (**Supplemental Figure S6B, C** and **D online**). Consistent with the increased sensitivity of the *opt3* mutant to copper deficiency and our finding that it experiences copper deficiency, cupric reductase activity was significantly higher in roots of the *opt3* mutant *vs.* wild type (**Supplemental Figure S6E online**).

**Figure 4.**
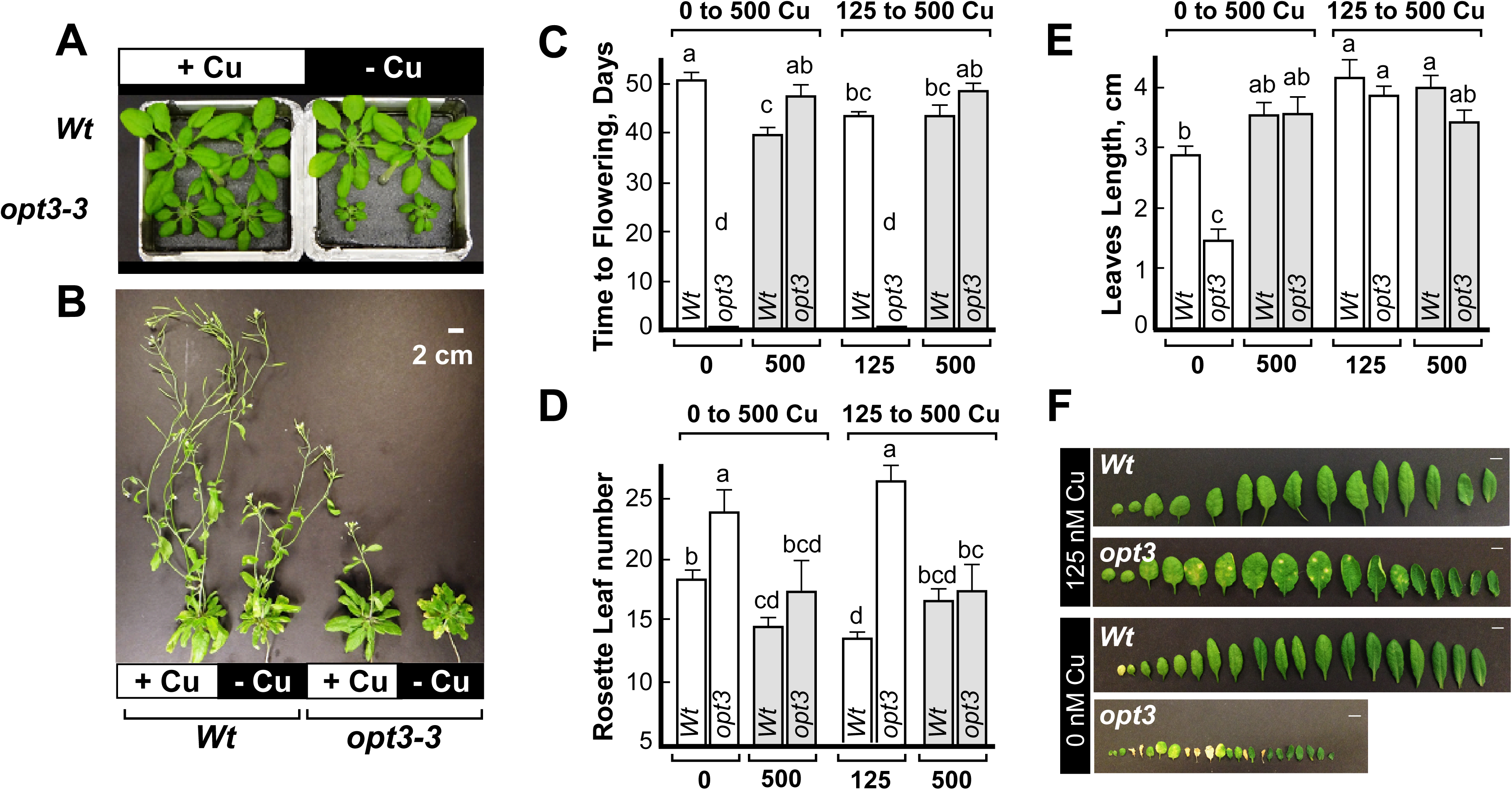
The *opt3* mutant is sensitive to copper deficiency. In (**A**) and (**B**), wild type and the *opt3* mutant were grown hydroponically with or without 125 nM CuSO_4_ (+ Cu or – Cu, respectively). Images were taken after four weeks (**A**) or nine weeks (B) from the seed sowing. (**C**), (**D**) and (**E**) show time to flowering, primary rosette leaf number and leaf length, respectively. Plants were germinated and grown hydroponically without or with copper (**0** or **125**, respectively and **white bars**). After four weeks of growth, a subset of plants grown with or without copper was transferred to 500 nM CuSO_4_ (**500** and **grey bars**). Measurements were taken after one week of growth and when plants were eight-week-old. Data show mean values ± S.E. Levels not connected by the same letter are statistically different (Tukey HSD, JMP Pro 14 software package, n = 3 to 6 measurements (plants) showing a representative outcome of at least three independent experiments). Data for the *opt3-2* allele *vs.* wild type are shown in **Supplemental Figure 6A**. (**F**) shows a representative image of leaves (from young to old in the direction from left to right) of wild type and the *opt3* mutant, both grown hydroponically for four weeks with or without copper (**125 nM Cu** or **0 nM Cu**, respectively). **A**, **B** and **F** show a representative image of plants from three independent experimental set-ups.

We then tested whether the transition to flowering is delayed in the *opt3* mutant under control conditions and/or under copper deficiency because we recently found that copper is involved in this process (Rahmati Ishka and Vatamaniuk, 2020). Consistent with our recent findings, wild type plants flowered significantly later and developed more rosette leaves when grown without *vs.* with copper (**Figure 4B** to **D** and **Table 1**). The *opt3* mutant failed to flower within the time frame of the experiment (8 weeks) and developed 30%- and 90% more rosette leaves than wild type in the medium without or with CuSO_4,_ respectively (**Figure 4C, D** and **Table 1**). Leaves of the *opt3* mutant were significantly shorter and were extensively chlorotic compared to wild type, both grown without added copper (**Figure 4E, F**). Although the length of the rosette leaves of mutant and wild type plants was comparable to control copper (125 nM CuSO_4_), the leaves of the mutant possessed characteristic chlorotic spots (**Figure 4F**). The delayed transition to flowering, the increased number, and the length of rosette leaves of the mutant were rescued by transferring the mutant to the medium with high (500 nM) CuSO_4_ (**Figure 4C to E, Table 1** and **Supplemental Figure 7A online**). Transferring the mutant to a lower (250 nM) copper also rescued the small size of the mutant, although to a lesser extent compared to the higher copper concentration (500 nM) (**Supplemental Figure 7B online**).

**Table 1.** Average day from the date of sawing to flowering in wild type and the *opt3-3* mutant grown hydroponically with 0 or 125 nM CuSO_4_. A subset of plants was shifted to 500 nM copper^1^.

| Cu concentrations, nM |  |  |  |  |
| --- | --- | --- | --- | --- |
| Genotype | 0 | 0 to 500 | 125 | 125 to 500 |
| Wild type (Col-0) | 46.83 ± 2.86 | 40.17 ± 0.79 | 42.33 ± 0.76 | 41.80 ± 1.24 |
| *opt3-3* (Col-0) | N.A.<sup>2</sup> | 48 ± 1.78 | N.A.<sup>2</sup> | 48.4 ± 1.80 |

<sup>1</sup> data showing average ± standard errors (n = 3 to 6)
<sup>2</sup> NA; not applicable. The mutant did not flower during the course of measurements which ranged from week 5 to week 8.

### The *opt3* Mutant Mounts Transcriptional Iron-Deficiency Responses in Roots but not in Shoots

We next used deep transcriptome sequencing to test whether the expression of copper deficiency-responsive genes is altered in roots, mature and young leaves of the *opt3* mutant compared to wild type. Using Illumina sequencing, we obtained 54, 79 and 59 million clean reads from roots, mature and young leaves, respectively (**Supplemental data set 1**). Of these, 86% of reads from roots and mature leaves and 93% of reads from young leaves were mapped to the *A. thaliana* genome and employed to estimate the transcript abundance and differential expression. Compared to wild type, 376, 673 and 1,942 genes were differentially expressed in the *opt3-3* mutant roots, young leaves and mature leaves, respectively (ratio ≥ 1.5 or ≤ 0.67, false-discovery rate [FDR] < 0.05; **Fig. 5A to C**). As expected, the expression of canonical iron deficiency-responsive genes, known to positively regulate iron-deficiency responses to facilitate iron uptake (*e.g., FIT*, *IRT1*, *FRO2*, *bHLH38*, *bHLH39*, *bHLH100*, *bHLH101, MYB10, MYB72),* iron-sensing peptide *FEP2/IMA2*, coumarin synthesis and transport (*CYP82C4*, *S8H* and *PDR9/ABCG37*) was highly up-regulated in roots of the *opt3* mutant (**Figure 5B** and **Supplemental Data Set 2**). In addition, IRT1 polypeptide was detected in roots of the *opt3* mutant grown under control conditions and roots of wild type grown under iron deficiency but not in roots of wild type or the *fit-2* mutant grown under control conditions (**Supplemental Figure S8 online**). Of other iron deficiency-regulated genes, the chloroplast-localized *FRO3* and iron exporter *IREG3/FPN3*, which is dual-targeted to mitochondria and chloroplast, were also upregulated in roots of the *opt3* mutant compared to the wild type (**Figure 5B** and **Supplemental Data Set 2)**. Of 237 genes upregulated in roots of the *opt3* mutant, 32 (13.5%) were among the robust FIT targets and 6 (0.025%) were among PYE targets (**Supplemental data set 2** and (Mai et al., 2015)), suggesting that the *opt3* mutant mounts primarily FIT-regulated iron deficiency response under iron sufficiency.

**Figure 5.**
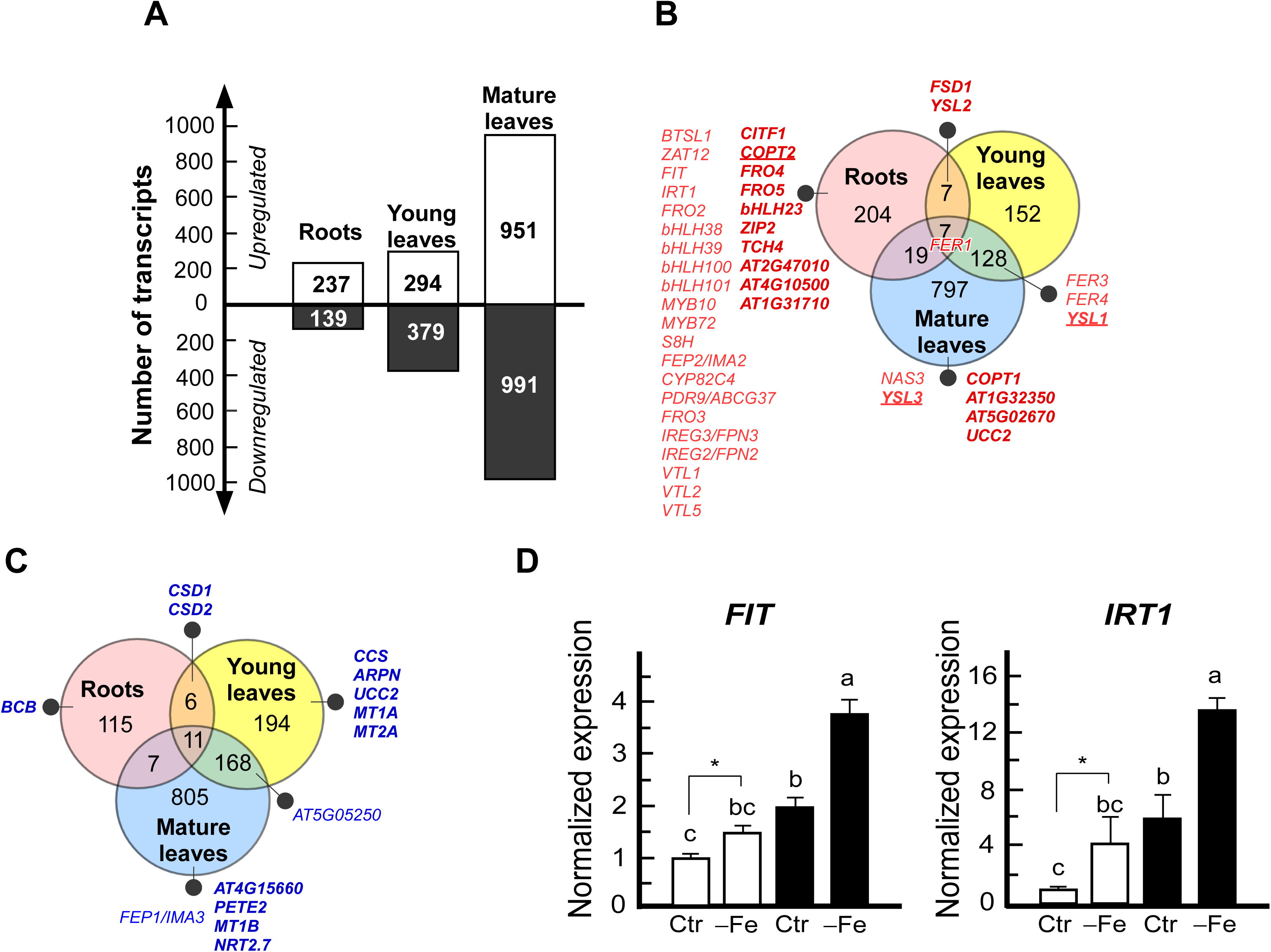
The *opt3* mutant mounts copper deficiency response in roots and young leaves. (**A**) Total number of differentially expressed genes in roots, mature leaves and young leaves of the *opt3* mutant *vs.* wild type, according to RNA-seq data (ratio >= 1.5 or <=0.67, false-discovery rate [FDR] < 0.05). Venn diagrams show the number of upregulated (**B**) or downregulated (**C**) genes in roots, mature and young leaves. Overlaps indicate the number of genes co-regulated in indicated tissues. Genes associated with iron and copper homeostasis are shown in (**B**) and (**C**). Genes involved in copper homeostasis are marked in bold; genes associated with iron and copper deficiency responses are marked in bold and underlined. (**D**) The expression of the iron transport system is upregulated in the roots of the *opt3* mutant under iron deficiency. Plants were grown hydroponically for 31 days. A subset of plants was transferred to a fresh medium without iron. Roots were collected after four days of additional growth with or without iron. Shown are mean values ± S.E. (n = 3 independent RT-qPCR experiments per genotype. In each experiment, roots were pooled from four plants grown in one container for RNA extraction). Asterisks indicate statistically significant differences in gene expression in wild type grown with or without iron (*p* < 0.05, Student-*t* test). Levels not connected by the same letter are statistically different (*p* < 0.05; Tukey HSD, JMP Pro 14 software package.

Notably, the expression of a negative FIT regulator, *ZAT12*, was upregulated in roots of the *opt3* mutant *vs.* wild type as well (**Figure 5B** and **Supplemental Data Set 2**). In addition, the expression of genes mediating cellular response to iron overload was upregulated. Specifically, the expression of *FER1* encoding a chloroplast-localized iron-sequestering protein, whose expression is upregulated by iron overload to protect chloroplasts from iron toxicity, was upregulated in roots of the *opt3* mutant *vs.* wild type (**Figure 5B** and **Supplemental Data Set 2)**. The expression of *IREG2/FPN2, VTL1, 2, 5* mediating iron sequestration into the vacuole was upregulated too (**Figure 5B** and **Supplemental Data Set 2)**. These data suggest that despite the upregulated FIT-transcriptional network, root cells also perceived the iron-sufficiency/overload signal and responded by increasing the expression of FIT negative regulators and genes involved in the mitigation of iron-overload toxicity. It is noteworthy that while roots of the *opt3* mutant constitutively overexpressed genes encoding iron uptake system even when grown in iron sufficient conditions, the magnitude of the response was significantly larger when mutant was grown under iron deficiency (**Figure 5D**). This is consistent with the important role of OPT3 in mitigating and controlling the basal response of the root to iron deficiency.

We then compared the expression of iron-deficiency responsive genes in young and mature leaves of the *opt3* mutant *vs.* wild type. Unlike roots, both mature and young leaves of the *opt3* mutant did not show transcriptional iron deficiency response. Specifically, none of the canonical iron-deficiency upregulated genes were upregulated in young or mature leaves of the *opt3* mutant (**Figure 5B** and **Supplemental Data Set 3 and 4**). Moreover, *FEP1/IMA3*, shown to be involved in iron sensing and typically upregulated in leaves and roots under iron deficiency (Grillet et al., 2018), was downregulated in mature leaves of the *opt3* mutant *vs.* wild type; *At5g05250*, encoding a protein with unknown function, was upregulated by iron deficiency in leaves of different *A. thaliana* ecotypes (Waters et al., 2012) was downregulated in both young and mature leaves of the *opt3* mutant *vs.* wild type. Iron deficiency downregulated genes *NAS3*, *YSL1* and *YSL3* were highly upregulated in mature leaves in the *opt3* mutant, and *YSL1* was also upregulated in young leaves of the *opt3* mutant compared to wild type (**Figure 5B** and **Supplemental Data Set 3 and 4**). The chloroplast-localized ferric chelate reductase *FRO7* and plasma membrane-localized *FRO6*, although not regulated by iron deficiency in shoots of *A. thaliana* (Mukherjee et al., 2006), were highly upregulated in mature leaves of the *opt3* mutant *vs.* wild type (**Figure 5B** and **Supplemental Data Set 3**). At the same time, iron-sufficiency markers, *FER1, FER3* and *FER4*, were highly upregulated in both mature and young leaves of the *opt3* mutant compared to wild type (**Figure 5B** and **Supplemental Data Set 3 and 4**). Together, our RNA-seq data are helping to unravel the details of contrasting responses of the iron-regulon in roots, young and mature leaves of the *opt3-3* mutant and supported the past observation that leaves of the *opt3* mutants sensed iron overload (Khan et al., 2018).

### The *opt3* Mutant Mounts Transcriptional Copper Deficiency Response

Roots of the *opt3* mutant accumulated significantly less copper than roots of wild type plants, albeit the total internal concentration in roots was at the sufficiency level (Fig. 2A and (Broadley et al., 2012)). We thus, anticipated that the expression of genes belonging to copper-deficiency regulon in roots would not be altered in the *opt3* mutant. However, we found that the expression of canonical copper-deficiency induced genes, responsible for copper uptake (*CITF1*, *COPT2*, *FRO4* and *FRO5*) was up-regulated in roots of the *opt3* mutant compared to wild type (**Table 2, Figure 5B** and **Supplemental Data Set 2**). Since small RNAs were not included in the RNA-seq analysis, we tested the expression of canonical copper miRNAs expression in roots of the *opt3* mutants *vs.* wild type plants by RT-qPCR (Pilon, 2017). We found that the expression of copper deficiency regulated micro RNAs, *miR397a/b*, *miR857a*, *miR389b/c* and *miR408* was upregulated in roots of the *opt3* mutant vs. wild type (**Figure 6A**). Also, copper deficiency-repressed genes, contributing to copper economy/metal switch, *CSD1*, *CSD2* and *BCB*, encoding Cu/Zn superoxide dismutases 1 and 2 and blue-copper-binding protein, respectively, were downregulated by more than 2-fold. Consistently, the expression of *FSD1* (Fe-containing superoxide dismutase) was upregulated by 2.7-fold in roots of the *opt3* mutant *vs.* wild type (**Table 2, Figure 5C** and **Supplemental Data Set 2**). In addition, several other canonical copper-deficiency upregulated genes (*bHLH23, ZIP2*, *YSL2, TCH4, AT2G47010, AT4G10500*, *AT1G31710*) were upregulated in roots of the *opt3* mutant compared to wild type (**Table 2, Figure 5B** and **Supplemental Data Set 2)**. Of 16 copper-deficiency regulated genes in the *opt3* mutant, 10 are regarded as SPL7-dependent (**Supplemental Data Set 2)**. These results show that roots of the *opt3* mutant manifest molecular symptoms of copper deficiency even though plants were grown under copper-sufficient conditions and the internal concentration of copper in roots was at the level of sufficiency (**Figure 2A** and **Supplemental Figure online 3**).

**Figure 6.**
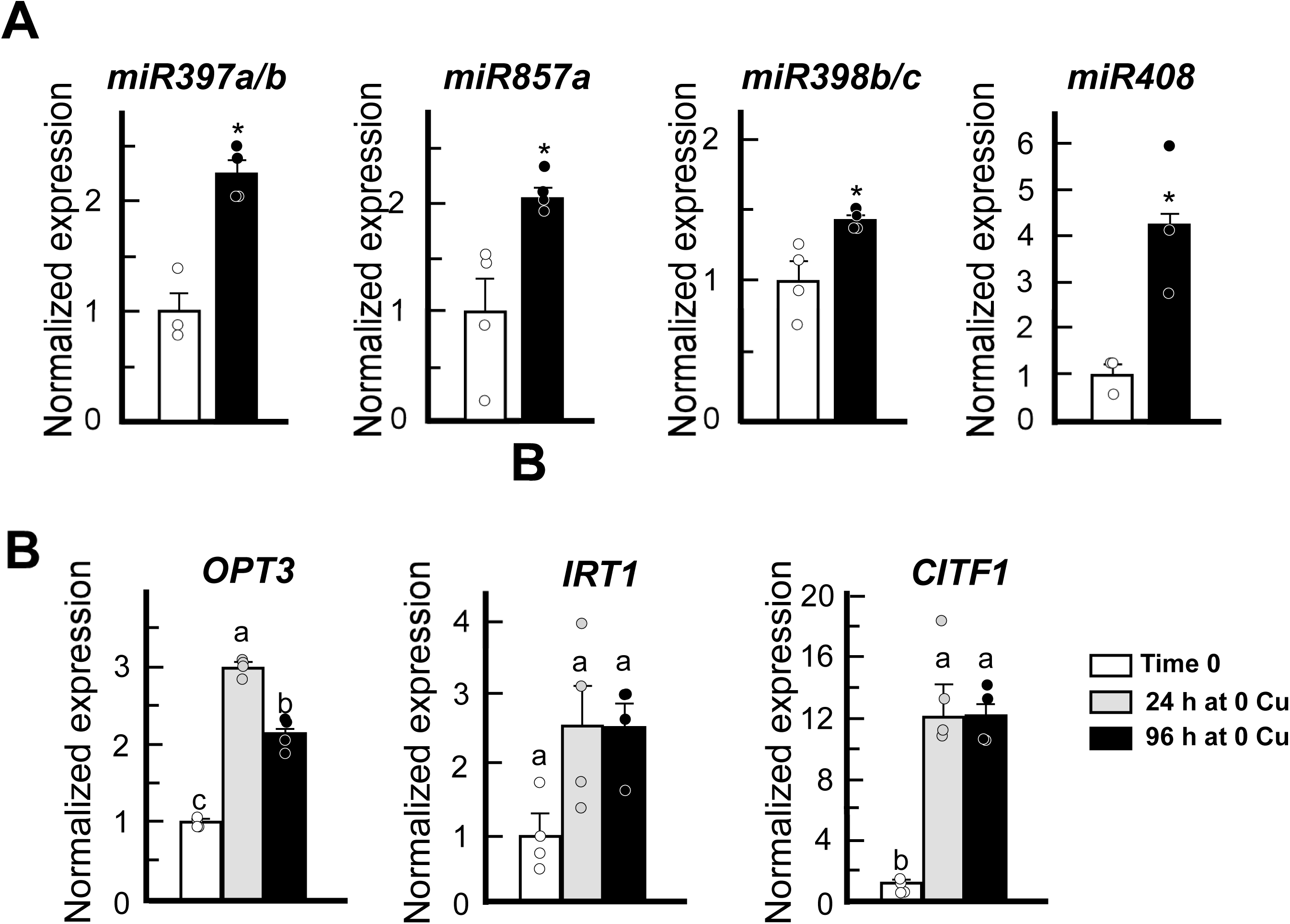
The expression of copper micro-RNA genes is upregulated in the *opt3* mutant. **(A)** shows the transcript abundance of *miRNA397a/b*, *miR857a*, *miR39b/c*, and *miR408* in roots of 5-week-old wild type (open bars) or the *opt3* mutant (black bars) grown hydroponically with 125 nM CuSO_4_. Shown are mean values ± S.E. Asterisks indicate statistically significant differences from the expression of genes in wild type, set to one (*p* < 0.05, Student-*t* test, n = 4 independent RT-qPCR experiments per genotype. In each experiment, roots were pooled from four plants grown in one container for RNA extraction). (**B**) Expression of *OPT3*, *IRT1* and *CITF1* in roots of wild type subjected to one or four days under copper deficiency. *CITF1* was used as a marker of copper deficiency to validate treatment efficiency. Shown are mean values ± S.E. Levels not connected by the same letter are statistically different (p < 0.05; Tukey HSD, JMP Pro 14 software package, n = 4 independent RT-qPCR experiments per condition. In each experiment, roots were pooled from four plants grown in one container for RNA extraction.

**Table 2.**
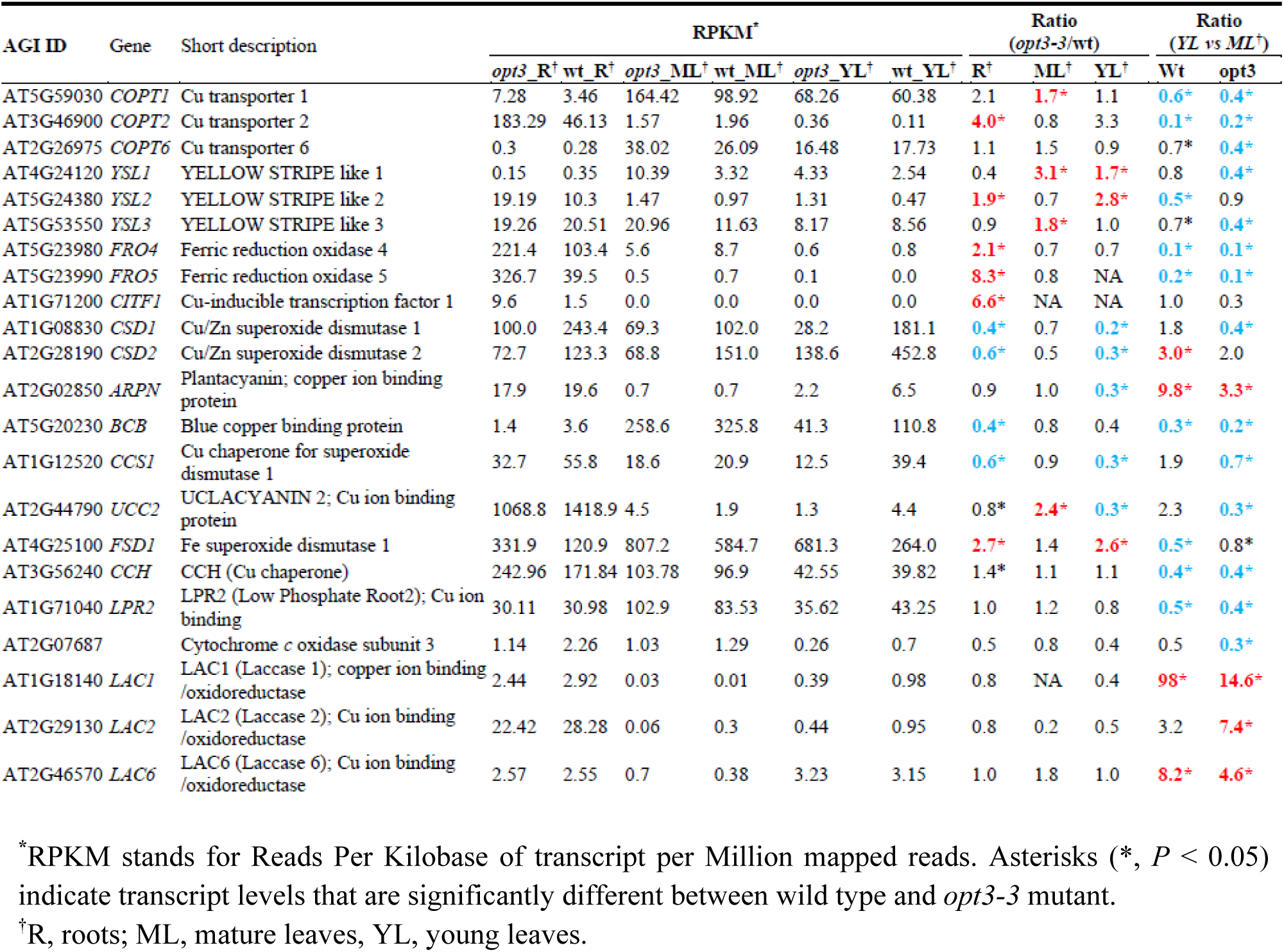
Expression of copper deficiency-responsive genes in roots, mature leaves and young leaves of the *opt3-3* mutant. Upregulated or downregulated genes are shown in red or blue bold font, respectively, and marked with asterisks (ratio >= 1.5 or <=0.67, [FDR] < 0.05).

We also found that young leaves of the *opt3* mutant manifested molecular symptoms of copper deficiency as evident by the increased expression of *YSL1* and *YSL2*, typically upregulated under copper deficiency and involved in lateral movement of minerals, including copper (**Figure 5B, C** and **Table 2, Supplemental Data Set 3**). Genes associated with either copper buffering *(MT1A* and *MT2A*) or copper economy (*CSD1*, *CSD2*, *CCS1*, *ARPN, UCC2)* were downregulated, while *FSD1* was upregulated (**Figure 5B, C** and **Supplemental Data Set 3**).

Regarding mature leaves, a different set of genes was differentially expressed in the *opt3* mutant *vs.* wild type, and the pattern of the regulation (up or down) of canonical copper-deficiency-regulated genes was not symptomatic for either deficiency or sufficiency (**Supplemental Data Set 4)**. Specifically, as would be expected under copper deficiency, the expression of genes associated with copper uptake and lateral movement, *COPT1*, *YSL1* and *YSL3,* was upregulated in the *opt3-3* mutant *vs.* wild type. Other *A. thaliana* copper-deficiency upregulated genes, including *AT1G32350* and *AT5G02670*, were also upregulated in mature leaves of the *opt3* mutant. Of genes typically downregulated by copper deficiency, *AT4G15660* was also downregulated in the *opt3* mutant *vs.* wild type as well as *PETE2*, associated with copper sparing and *MT1B*, associated with copper buffering. By contrast, the expression of another copper sink, *UCC2*, typically downregulated by copper deficiency, was upregulated in mature leaves of the *opt3* mutant *vs.* wild type while *NRT2.7*, typically upregulated by copper deficiency in *A. thaliana*, was downregulated in the *opt3* mutant *vs.* wild type.

To conclude, our RNA-Seq data show that both roots and young leaves of the *opt3* mutant mounted transcriptional copper deficiency response while only roots of the mutant manifested transcriptional iron deficiency response. In addition, our finding of the distinct transcriptional response and metal accumulation in mature *vs.* young leaves of the mutant emphasizes the need to separate these leaves in analyses of mutant phenotypes.

### *OPT3* is Transcriptionally Upregulated by the Short-term Copper Deficiency

While *OPT3* is robustly upregulated in roots and leaves by iron deficiency, it was not found among copper-deficiency-responsive genes in the existing RNA-seq data. In these RNA-seq analyses, plants were exposed to copper deficiency for a minimum of three days. Since *OPT3* responds to iron deficiency within 24 h (Khan et al., 2018), we hypothesized that *OPT3* might also be transcriptionally upregulated by short-term exposure to copper deficiency. Thus, we compared the *OPT3* expression in *A. thaliana* subjected to copper deficiency for 24 or 96 h. We also examined the expression of a copper-deficiency marker, *CITF1,* to validate the efficiency of treatments. *CITF1* was up-regulated in roots by 16-fold after 24 h of copper deficiency and remained highly upregulated after 96 h. The expression of *OPT3* was also upregulated, although to a lesser extent, after 24 h of copper deficiency, but unlike *CITF1*, the transcript abundance of *OPT3* decreased after 96 h of treatment (**Figure 6B**). The expression of *IRT1* was not upregulated by copper deficiency (**Figure 6B**).

### The Molecular Symptoms of Copper Deficiency in the *opt3* Mutant are Rescued by Supplemental Copper

Since transferring the *opt3* mutant to higher copper concentrations decreased its time to flowering to the level of wild type and rescued the length of rosette leaves (**Figure 4** and **Supplemental Fig. 7A online**), we predicted that supplemental copper would also decrease the expression of copper-deficiency responsive genes. To test this hypothesis, we compared the transcript abundance of copper deficiency markers, *CITF1*, *COPT2*, *FRO4* and *FRO5* in roots of the *opt3* vs. wild type. In parallel, we also tested the expression of key iron deficiency markers, *IRT1* and *FRO2*. Consistent with RNA-seq data (**Table 2**), the expression of *CITF1*, *COPT2*, *FRO4* and *FRO5* was upregulated in roots of the *opt3* mutant *vs.* wild type, both grown under control conditions (**Figure 7)**. The transcript abundance of copper deficiency markers decreased in roots of the *opt3* mutant after transfer to higher concentrations of copper (**Figure 7A** to **H**). We note that transferring wild type plants to higher copper also decreased the expression of *CITF1, COPT2*, *FRO4* and *FRO5,* suggesting that 125 nM CuSO_4_ was somewhat copper limiting even though the growth and development of wild type plants were not affected. Supplemental copper in both concentrations also decreased the expression of *IRT1* and *FRO2* in roots of the *opt3* mutant compared to their expression levels under control conditions (**Figure 7 J to M**). It is noteworthy that the higher copper concentration (500 nM) increased the expression of both *IRT1* and *FRO2* in roots of wild type, reinforcing the existence of interactions between copper and iron homeostasis.

**Figure 7.**
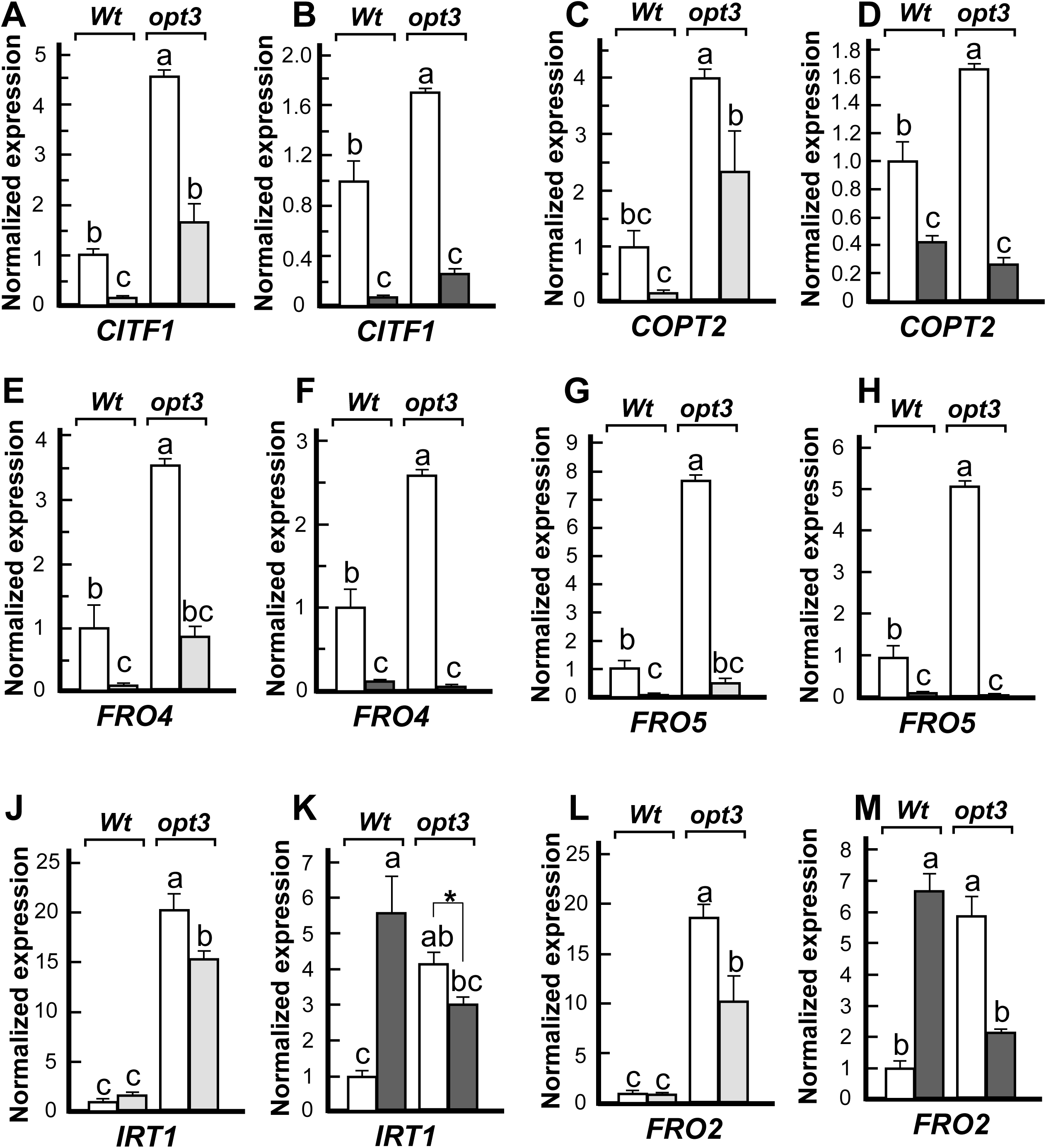
Copper supplementation partially rescues the molecular symptoms of copper deficiency in roots of the *opt3* mutant. Plants were grown hydroponically under control conditions (**white bars**) until the late-vegetative stage before a subset of plants was transferred to a fresh medium with higher copper concentrations (250 nM CuSO_4_, **light grey bars** in **A, B, E, F, J, K** or 500 nM CuSO_4_, **dark grey bars** in **C, D, G, H, L, M**). Plants were grown for another week before tissue collection and RT-qPCR analysis. The transcript abundance of indicated genes was normalized to the wild type grown under control conditions. Data show mean values ± S.E. Levels not connected by the same letter are statistically different (Tukey HSD, JMP Pro 14 software package, n= 3 RT-qPCR experiments per genotype. In each experiment, roots were pooled from three plants grown in one container for RNA extraction. Data are normalized to the expression of *Actin2*; the expression of genes in wild type grown under 125 nM CuSO_4_ (control conditions) was set to one.

### Copper Deficiency Response in Roots of the *opt3* Mutant is Regulated by the Shoot

We next tested whether the increased expression of copper deficiency markers in roots of the *opt3* mutant is due to its altered shoot-to-root signaling. To do so, we used reciprocal grafting with wild type and *opt3* plants and examined the transcript abundance of copper deficiency markers *CITF1*, *COPT2*, *FRO4*, *FRO5* in roots (**Figure 8)**. The expression of these genes was up-regulated in roots of grafted *opt3*/*opt3* (*opt3* scions grafted to *opt3* rootstocks) compared to the grafted WT/WT (wild type scions grafted to wild type rootstocks [control grafts], **Figure 8A**). The expression of *CITF1*, *COPT2*, *FRO4* and *FRO5* was also elevated in roots of grafts with *opt3* scions and wild type rootstock (**Figure 8A**). In contrast, grafting of the wild type shoots onto the *opt3* rootstocks downregulated *CITF1*, *COPT2*, *FRO4* and *FRO5* expression relative to control grafts (**Figure 8A**). These data show that the OPT3 function in the shoot is sufficient to regulate the transcriptional copper deficiency responses in the root.

**Figure 8.**
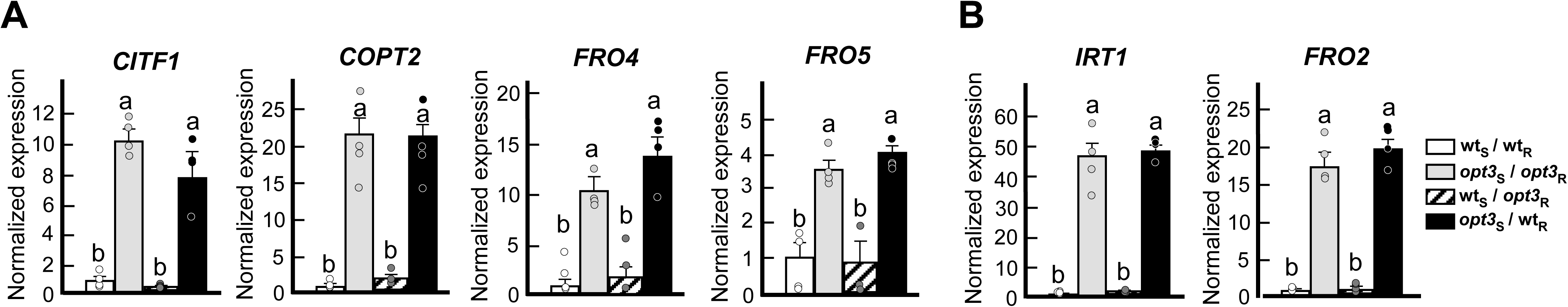
The OPT3 function in the shoot controls the expression of copper deficiency markers. The transcript abundance of copper- (**A**) and iron-deficiency (**B**) markers in roots of grafted plants. Wild type and the *opt3* mutant were used for reciprocal grafting. **Wt*S*/Wt*R*,** wild type scion grafted to wild type rootstock (control); ***opt3S/opt3R***, *opt3* scion grafted to *opt3* rootstock; **Wt*S*/opt3*R***, wild type scion grafted to *opt3* rootstock; ***opt3S/WtR***, *opt3* scion grafted to wild type rootstock. Shown values represent means ± S.E. (n = 4 independent experiments [PCR runs, each representing an independent experimental setup]; roots were pooled from four plants per experimental set-up). Levels not connected by the same letter are statistically different (*p* < 0.05; Tukey HSD, JMP Pro 14 software package). Data are normalized to the expression of *Actin2*; the expression of genes in wild type was set to one.

Grafted plants performed as expected, as evidenced by the expression of the iron-deficiency markers, *IRT1* and *FRO2* in different graft combinations (**Figure 8B** and (Zhai et al., 2014)). As we showed previously, *IRT1* and *FRO2* were up-regulated in roots of *opt3*/*opt3* compared to WT/WT grafts (**Figure 8B** and (Zhai et al., 2014)). Grafting wild type shoots onto *opt3* mutant roots downregulated *IRT1* and *FRO2* expression relative to their expression in control grafts (**Figure 8B**). In contrast, grafting of *opt3* shoots onto wild type roots increased the expression of iron deficiency markers (**Figure 8B**). Together, these data show that OPT3 function in the shoot regulates both iron and copper deficiency responses of the root. These data also suggested the existence of the systemic shoot-to-root copper status signaling.

### Copper Movement *via* the Phloem from the Shoot to the Root Regulates the Expression of Copper-deficiency Responsive Genes

To test whether copper status of the shoot can be communicated to the root, and whether copper movement *via* the phloem from the shoot to the root influences the expression of copper acquisition genes, we performed phloem feeding experiments *via* the abrasion of the leaf vasculature. We first tested this approach by using the phloem symplastic tracer 5,6 carboxyfluorescein-diacetate (CFDA) (Grignon et al., 1989; Oparka et al., 1994). After diffusing into plant cells, CFDA is cleaved by cellular esterases to yield a membrane-impermeable form of the dye, carboxyfluorescein (CF). As expected, we detected CF accumulation in roots of wild-type plants after CFDA solution feeding *via* the incision in the leaf vasculature (**Supplemental Figure 9A, B online**).

Since systemic signaling of iron deficiency has already been established, we first exposed *Arabidopsis* wild type to iron deficiency and tested whether iron feeding to the phloem in the shoot would rescue the expression of iron deficiency markers *FIT* and *IRT1.* Since *COPT2* is also upregulated by iron deficiency *via FIT*, we also used *COPT2* as a marker for iron as well as copper deficiency. Consistent with the role of iron in the phloem in systemic signaling, feeding iron directly to the phloem of iron deficient plants decreased the expression of *FIT*, *IRT1* and *COPT2* (**Figure 9A**). We also noticed that iron but not mock-feeding, rescued the chlorosis of young leaves of iron-deficient plants (**Supplemental Figure 9D online**).

**Figure 9.**
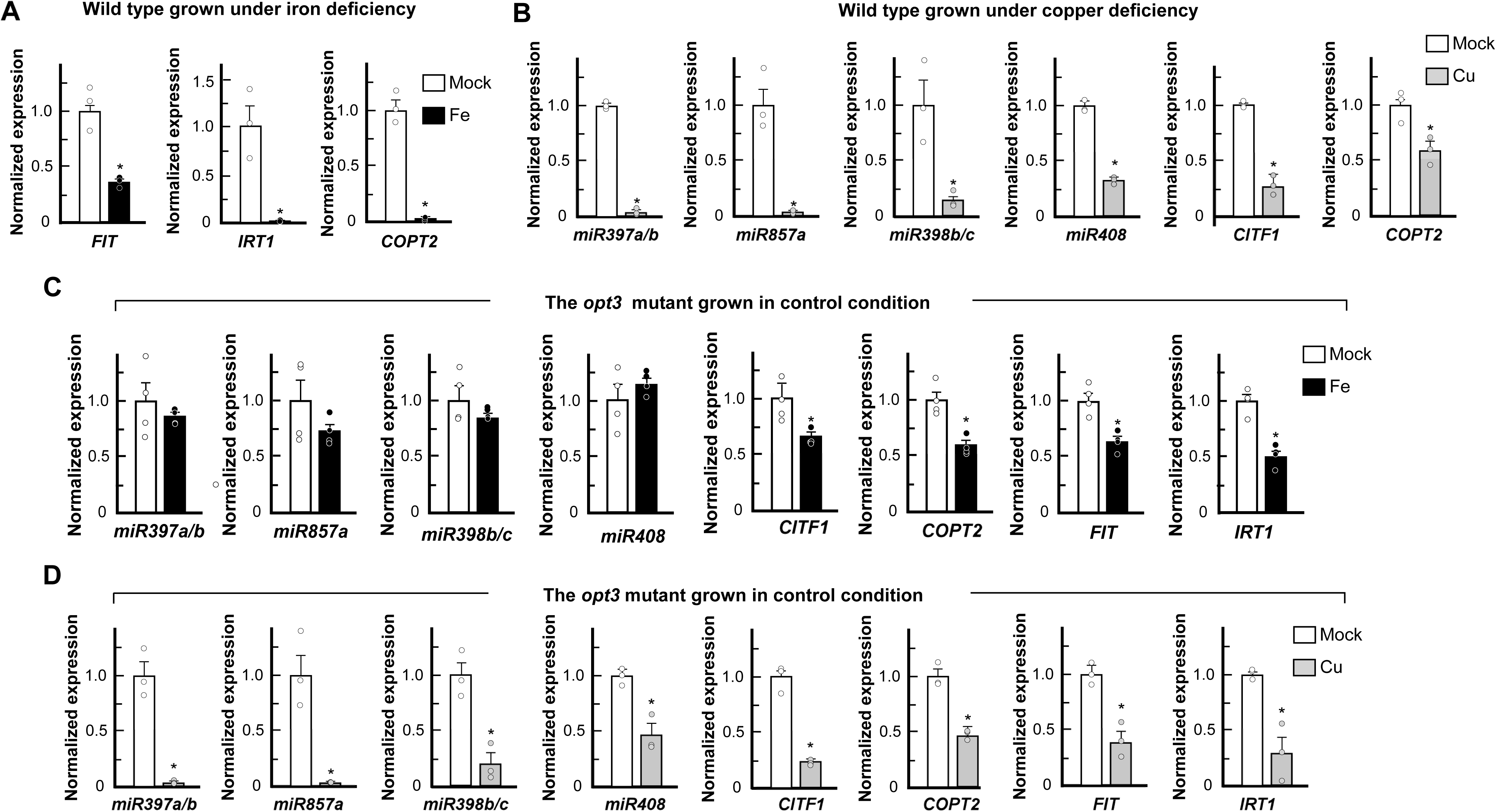
Copper or iron feeding *via* the phloem in the shoot represses the molecular symptoms of copper and iron deficiency in the root of the wild type and the *opt3* mutant. RT-qPCR analysis of copper and iron deficiency molecular markers in roots of the 33-day-old wild type (**A, B**) or the *opt3* mutant (**C, D**). Wild type plants were grown hydroponically and subjected to either iron or copper deficiency (**A** and **B**, respectively). The *opt3* mutant was grown hydroponically in copper and iron sufficient conditions (**C**, **D**). Roots were collected after 24h of phloem feeding with iron (**Fe**, black bars) or copper (**Cu**, grey bars) or control feeding (**mock**, open bars). Mock treatment for copper feeding was MiliQ water, and mock treatment for iron feeding was citrate. Asterisks indicate statistically significant differences from the expression of genes in mock-fed plants, set to one (*p* < 0.05, Student-*t* test, n = 4 [PCR runs, each representing an independent experimental set up]; roots were pooled from four plants per each experimental set-up). Data are normalized to the expression of *Actin2*; the expression of genes in a mock-treated wild type (**A**, **B**) or a mock-treated *opt3* mutant (**C**, **D**) was set to one.

We then exposed wild type to copper deficiency, fed copper to the phloem in the shoot and found that the expression of the copper deficiency markers, *CITF1*, *COPT2* and copper miRNAs, *miR397a/b*, *miR857a, miR389a/b* and *miR408* was significantly downregulated in the root (**Figure 9B**). These data show the existence of systemic signaling of copper deficiency in *A. thaliana* and the important role of copper movement *via* the phloem from the shoot to the root in the regulation of copper-responsive genes in the root. Our data also provide evidence that iron in the leaf phloem plays a repressive role in regulating iron deficiency-responsive genes.

### Feeding Copper or Iron to the Phloem in the Shoot of the *opt3* Mutant Attenuates the Expression of both Iron- and Copper-deficiency Responsive Genes in the *opt3* Mutant Roots

Past studies of foliar iron application by iron spraying did not rescue the expression of iron acquisition genes, constitutively overexpressed in roots of the *opt3* mutant (Garcia et al., 2013). This finding is not surprising considering the *opt3* mutant’s defect in iron and copper loading to the phloem in leaves (**Figures. 1F, G, K** and **5B** and (Zhai et al., 2014). We, thus, asked whether feeding with iron *via* the vasculature in the shoot would rescue the expression of iron deficiency markers, *FIT* and *IRT1* in the root. Consistent with the suggested repressive role of iron movement *via* the phloem from the shoot to the root, the expression of *FIT* and *IRT1* was downregulated in roots of iron- *vs.* mock-fed plants (**Figure 9C**). Unexpectedly, we also found that the expression of copper deficiency markers, *CITF1* and *COPT2* was downregulated in the root of iron *vs.* mock-fed mutant (**Figure 9C**). Interestingly, the expression of copper miRNAs, *miR397a/b*, *miR857a, miR389a/b* and *miR408* in the root of the *opt3* mutant did not respond to the phloem feeding with iron in the shoot (**Figure 9C**).

Next, we fed copper to the phloem in the shoot to test whether it would have a similar repressive effect on the expression of iron and copper-deficiency regulated genes in roots of the *opt3* mutant. The expression of copper-deficiency markers including *CITF1*, *COPT2* and copper miRNAs, *miR397a/b*, *miR857a, miR389a/b* and *miR408* was significantly downregulated in the roots of copper *vs.* mock-fed mutant (**Figure 9D**). The expression of iron deficiency marker genes, *FIT* and *IRT1*, constitutively upregulated in the *opt3* mutant, was also downregulated in roots of the *opt3* mutant, fed with copper *via* the phloem in the shoot (**Figure 9D**). Together, these data suggest the existence of crosstalk between iron and copper in systemic signaling directed on the regulation of copper and iron uptake system.

### Phloem Copper or Iron Feeding Downregulates the Autonomously Upregulated Expression of *CITF1, copper miRNAs* and *FIT* in Roots of *A. thaliana* Under Simultaneous Copper and Iron Deficiency

We showed that copper and iron accumulation in the phloem was lower in the *opt3* mutant than in wild type and that OPT3 function is important for regulating copper and iron deficiency responses in the root (**Figures 1, 8** and **9**). To mimic the effect of simultaneous copper and iron deficiency in the phloem of the *opt3* mutant and its effect on the expression of corresponding deficiency markers in roots, we subjected *A. thaliana* wild type to deficiency of these metals applied concurrently. We tested the expression of *CITF1*, *miR397a/b*, *miR857a, miR389a/b* and *miR408* because these genes are highly regulated by copper deficiency and are downregulated by iron deficiency (Waters et al., 2012; Yan et al., 2017). We also tested the expression of *FIT* as it is responsive to iron but not copper deficiency (Colangelo and Guerinot, 2004; Bernal et al., 2012). As expected, copper and iron deficiency applied individually had an opposite effect on the expression of *CITF1, miR397a/b*, *miR857a, miR389a/b* and *miR408* (**Figure 10A**); *FIT* expression was upregulated under iron but not copper deficiency (**Figure 10A**). In contrast, all tested genes were highly upregulated when iron and copper deficiencies were applied simultaneously (**Figure 10A**). This result suggested that iron and copper deficiency applied simultaneously act in parallel/autonomously on the expression of copper and iron-responsive genes in the root.

**Figure 10.**
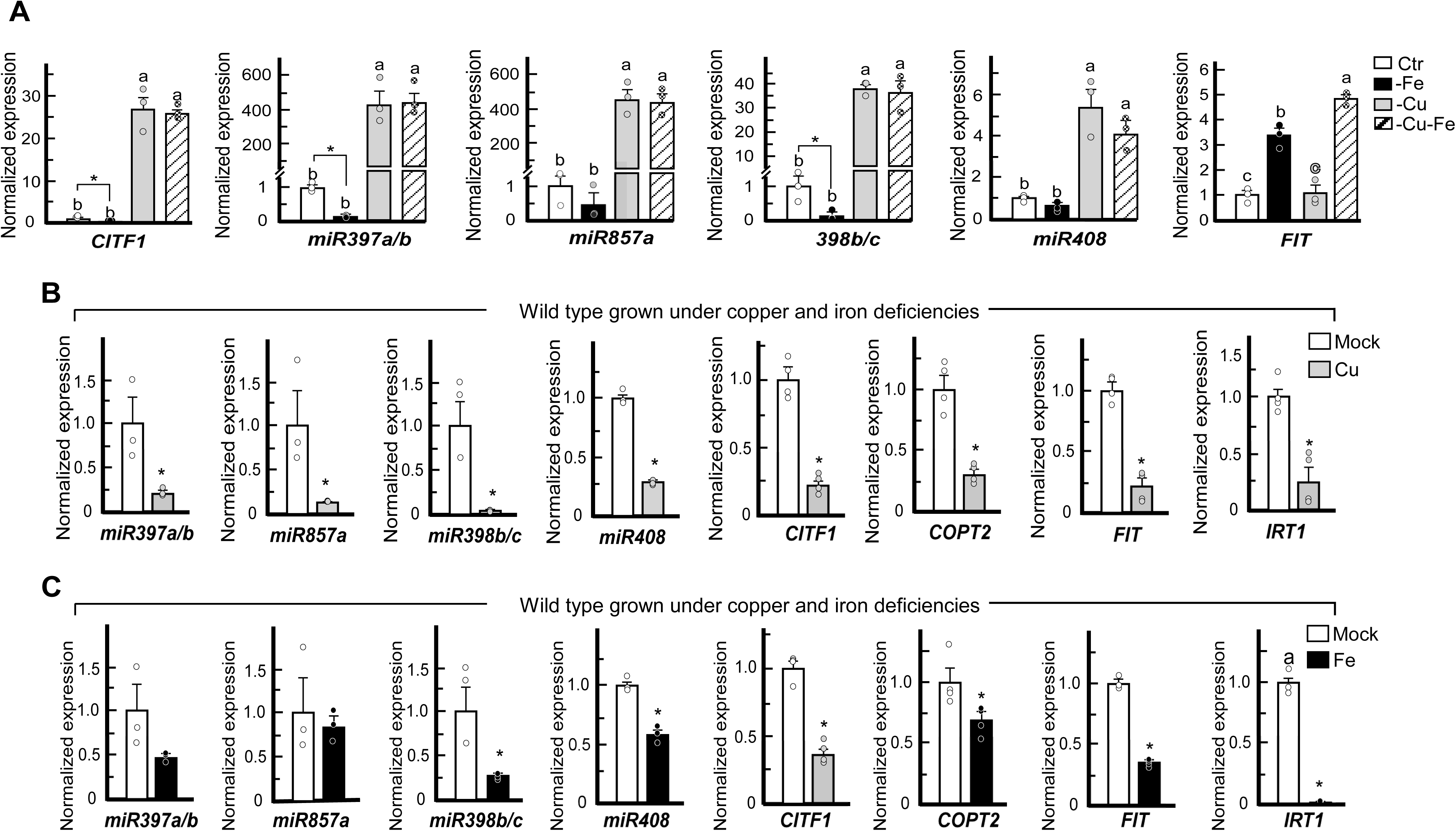
Copper and iron deficiency applied simultaneously increases the expression of *CITF1*, *copper miRNAs* and *FIT*. (**A**) Transcript abundance of *CITF1* and indicated *copper miRNAs* and the *FIT* was analyzed in roots of plants grown under control conditions (**Ctr**) or without cooper but with iron (**–Cu**) for four weeks before tissue collection. To achieve iron deficiency, plants were grown under control conditions for three weeks, transferred to a solution without iron and grown for one additional week (**–Fe**). For the simultaneous iron and copper deficiency treatment, plants were grown without copper but with iron for three weeks and then transferred to a fresh hydroponic medium lacking both copper and iron. Tissues were collected and analyzed after an additional week of growth (**–Cu –Fe**). (**B**) Transcript abundance of indicated genes in roots of wild type grown under simultaneous copper and iron deficiency after phloem feeding with copper (**Cu**, grey bars) or control feeding (**mock**, open bars). (**C**) Transcript abundance of indicated genes in roots of wild type grown under simultaneous copper and iron deficiency after phloem feeding with iron (**Fe**, black bars or control feeding (**mock**, open bars). Mock treatment for copper feeding was MiliQ water, and mock treatment for iron feeding was citrate. In (**A**), (**B**) and (**C**), mean values ± S.E are shown. Levels not connected by the same letter are statistically different (Tukey HSD, JMP Pro 14 software package, n = 3 to 4 independent experiments (PCR runs, representing an independent experimental set-up where roots were pooled from three plants per experiment). Data are normalized to the expression of *Actin2*; the expression of genes in wild type grown under control conditions (**A**) or mock-treated wild type (**B, C**) was set to one.

Copper feeding *via* the phloem in the shoot downregulated the expression of not only *copper-miRNAs* and *CITF1* but also *FIT* in the root of plants grown under simultaneous copper and iron deficiency (**Figure 10B**). Feeding with iron also downregulated not only the expression of *FIT* but also *CITF1*; of copper miRNAs, the expression of all, but not *miR857a,* was downregulated in roots of wild type grown under simultaneous copper and iron deficiencies (**Figure 10C**). Together, these data suggest that copper and iron in the phloem can mimic each other’s function in long-distance signaling function.

### Copper and Iron Deficiencies Applied Simultaneously Increase Root Iron Uptake and Delivery to Shoots While High Iron Reduces Copper Uptake

Because copper deficiency stimulates iron uptake, we hypothesized that copper deficiency in the *opt3* mutant and its lower copper and iron accumulation in the phloem compared to wild type, contributes to increased iron accumulation in its roots and leaves. To test this hypothesis, we exposed wild type to copper and iron deficiency individually and concurrently. We then analyzed iron accumulation in roots and leaves of wild type plants. As was found previously, copper deficiency increased iron accumulation in both roots and leaves of *A. thaliana* wild type compared to control conditions (**Figure 11A** and (Bernal et al., 2012; Waters et al., 2012; Kastoori Ramamurthy et al., 2018; Sheng et al., 2021)). As expected, iron accumulation was reduced in both roots and shoots under iron deficiency. Interestingly, the simultaneous application of copper and iron deficiency increased iron accumulation in both roots and shoots compared to iron deficiency applied individually (**Figure 11A**). We note that plants were subjected to iron deficiency for one week and prior to that they were grown under iron replete conditions. Thus, it is possible that root apoplastic iron reserves were absorbed and accounted for iron accumulation in shoots under simultaneous iron and copper deficiencies. It is also possible that simultaneous application of both copper and iron deficiency stimulated alternative high-affinity iron uptake pathways that contributed to the root absorption of residual iron in medium.

**Figure 11.**
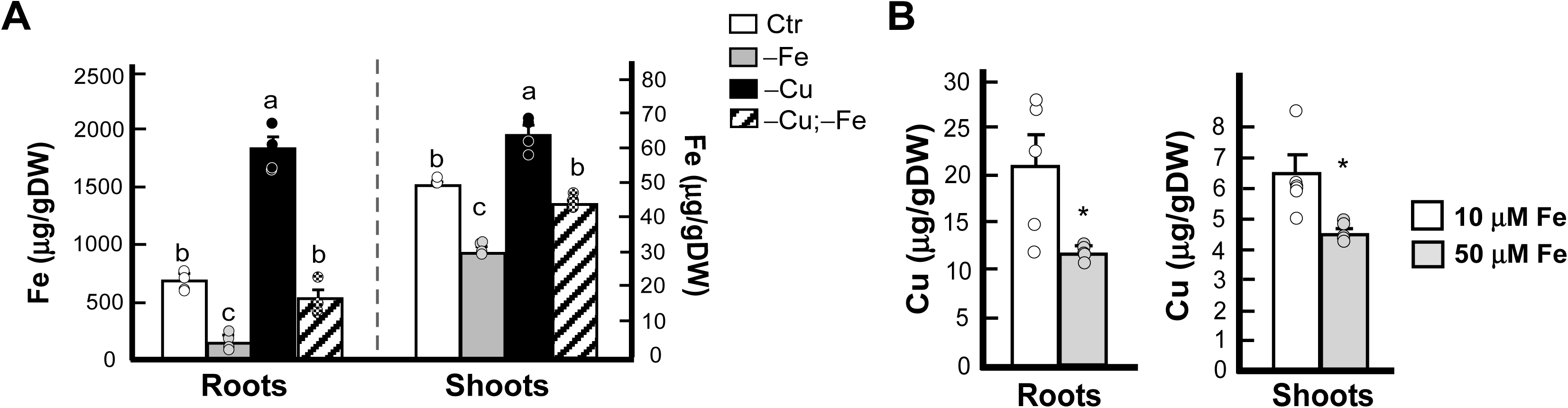
Copper and iron availability in the medium influence each other’s uptake and accumulation in roots and shoot of *A. thaliana*. (A) Iron concentration in roots and shoots of *A. thaliana* grown for five weeks in the control condition (**Ctr**), copper deficiency (**−Cu**), iron deficiency (**−Fe**), and simultaneous copper and iron deficiency (**−Cu −Fe**). Data are mean values ± S.E. Levels not connected by the same letter are statistically different (Tukey HSD, JMP Pro 14 software package, n = 3 to 5 independent ICP-MS measurements, each representing an independent experimental set-up; roots and shoots were pooled from four plants per each experimental set-up to obtain one measurement. (**B**) Copper concentration in roots and shoots of *A. thaliana,* grown for five weeks in the control condition (**10 µM Fe)** or with high iron (**50 µM Fe**). Mean values ± S.E is obtained with n = 5 ICP-MS measurements from two batches of independently grown plants per condition. Tissues from four plants were pooled for each measurement. Asterisks indicate statistically significant differences (*p* < 0.05, Student-*t* test).

Here, we also asked whether increased accumulation of iron in roots of the *opt3* mutant contributed to the reduced copper accumulation in its roots (**Figure 2A**). To mimic high iron conditions, we grew wild type plants under control (10 µM) or high (50 µM) iron in the medium and evaluated the accumulation of copper in its tissues. As expected, due to the toxic nature of iron, high iron in the medium reduced the root growth while increasing root iron level in the wild type compared to controls (**Supplemental Figure 10A, B online**). Iron accumulation in the root was accompanied with significantly reduced copper accumulation in roots and shoots (**Figure 11B**).

We recognize that these experiments do not perfectly match the scenario in the *opt3* mutant that experiences the internal copper and iron deficiency under control conditions. Nevertheless, our results favor the intriguing suggestion that decreased copper accumulation in roots and leaves and the reduced iron and copper accumulation in the phloem contribute to iron overaccumulation in the *opt3* mutant. We also propose that high iron in roots of the *opt3* mutant negatively effects copper accumulation in roots of the mutant compared to wild type.

## Discussion

Iron and copper are essential micronutrients and yet if their accumulation is not highly regulated, they are toxic for the growth and development of all organisms, including plants, and homeostasis of these elements is intricately related. The hallmark of this relationship is the increased accumulation of copper in plant tissues under iron deficiency and the overaccumulation of iron under copper deficiency (**Figure 11** and (Bernal et al., 2012; Waters and Armbrust, 2013; Kastoori Ramamurthy et al., 2018; Rai et al., 2021; Sheng et al., 2021). The toxicity of both metals is avoided by the ability of plants to regulate copper and iron transport systems in response to the fluctuation of copper and iron availability in the immediate root environment and the demands of the developing shoot. The *A. thaliana* phloem companion cells-specific iron transporter, OPT3, is involved in both signaling iron demand from shoots to roots and in transporting iron to developing tissues such as the seed. The systemic control of copper uptake into roots and its relationship to the systemic control of iron uptake in plants has not yet been documented. In this manuscript, we show the existence of systemic signaling of copper status, its relationship with systemic iron signaling, and the central role of AtOPT3 in these processes (**Figure 12**).

**Figure 12.**
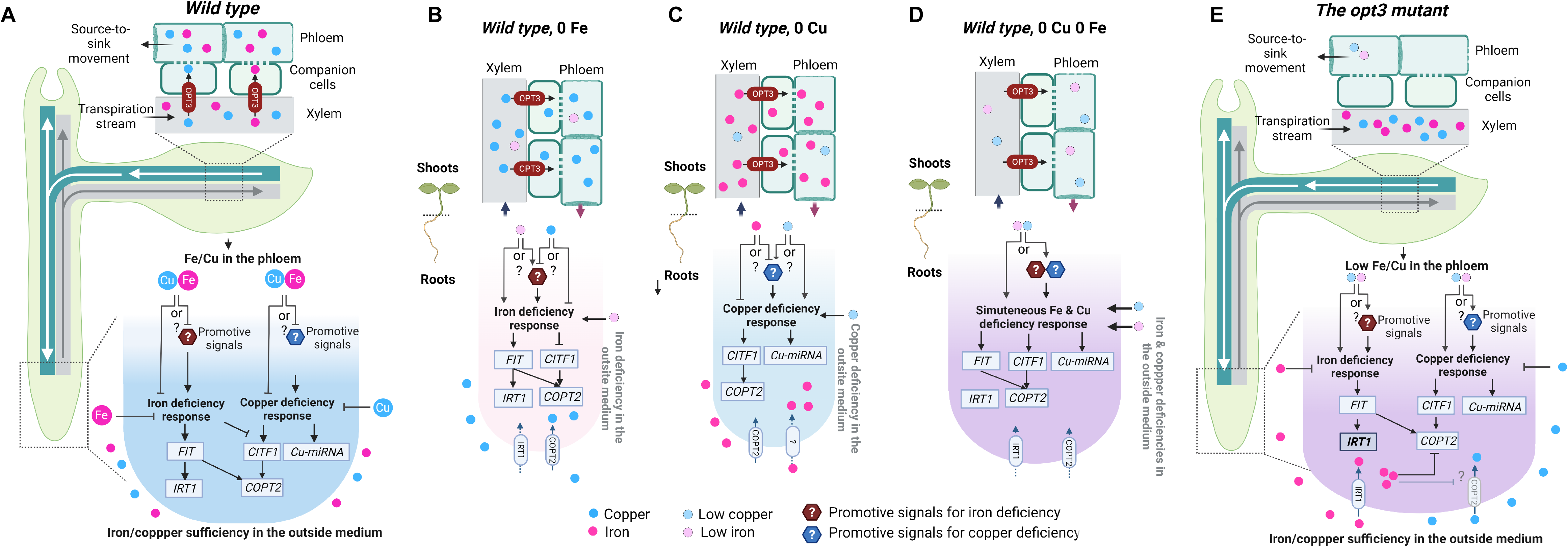
AtOPT3-facilitated Copper and Iron Transport to the Phloem Companion Cells Mediates Iron-Copper Crosstalk in Shoot-to-Root Signaling. (A) AtOPT3 mediates iron and copper loading to the phloem companion cells for the distribution to sink organs, including roots. Phloem-mobile iron and/or copper represses the expression of iron and copper uptake systems, either acting directly as a repressor or indirectly by sequestering or inhibiting the synthesis of a promotive signal. (**B**) Iron deficiency in the outside medium stimulates copper accumulation in roots and leaves (Waters et al., 2012; Waters and Armbrust, 2013; Waters et al., 2014) and **Figure 11A**. Under these circumstances, AtOPT3 loads mainly copper to the phloem in the shoot. The concentration of iron in the phloem decreases, which causes the upregulation of the expression of the iron uptake system. Copper availability in the phloem, provided by the AtOPT3, fine tunes the expression of iron transport genes to avoid iron overload observed in the *opt3* mutant. The repressive role of copper on the expression of copper and iron uptake system can occur directly or indirectly *via* scavenging or inhibiting the production of a promotive signal. A similar scenario occurs under copper deficiency, where OPT3 mainly loads iron to fine-tune the expression of copper uptake genes (**C**). Simultaneous iron and copper deficiencies in the outside medium autonomously upregulate the expression of both copper and iron uptake genes as both elements are less available in the phloem sap (**D**). The scenario is different in the *opt3* mutant (**E**): although both iron and copper are available in the outside medium, the *opt3* mutant experiences simultaneous iron and copper deficiency in the phloem. Low iron accumulation in the phloem stimulates transcriptional iron deficiency response, overriding local iron sufficiency response. Since iron is available in the outside medium, iron accumulates in the *opt3* mutant. Copper deficiency in the phloem, roots and leaves also leads to iron accumulation and transcriptional copper deficiency response. Increased iron accumulation from both pathways, in turn, decreases copper uptake, as shown in (**Figure 11B**) and documented in animal species (Klevay, 2001; Ha et al., 2016), further altering Fe-Cu crosstalk. Figure created in Biorender.com

### AtOPT3 Transports Copper and the Copper-NA Complex, but Free Copper Ions are Preferred Substrates in Heterologous Systems

The nature of the transport substrate of OPT3 has been under scrutiny and a matter of debate in the literature. OPT3 belongs to the OPT family that is phylogenetically divided into two groups: the Yellow Stripe-Like (YSL) and Oligopeptide Transporter (PT) clades (Lubkowitz, 2011). Analyses of the transport capabilities of some members of *A. thaliana* and *Oryza sativa* OPT family have disclosed their capacity to transport synthetic tetra- and pentapeptides in heterologous systems (Osawa et al., 2006), while members of the YSL clade are expected to transport metal-chelate complexes (Osawa et al., 2006). Given the role of OPT3 in iron homeostasis and signaling, it was expected to transport either peptides that can serve as metal ligands (*e.g.,* glutathione [GSH]) or metal-ligand complexes (*e.g.,* metal-nicotianamine [metal-NA]) (Stacey et al., 2008; Lubkowitz, 2011). Thus, our past finding that OPT3 transports free ions, iron and cadmium when expressed in *Xenopus* oocytes or the yeast iron uptake mutant, *fet3 fet4* was unexpected (Zhai et al., 2014). However, consistent with the proton-coupled nature of OPT-mediated transport, our past studies have shown that the acidification of uptake medium stimulated the OPT3-mediated inward currents (Osawa et al., 2006; Zhai et al., 2014). Past studies of the AtOPT3-GFP, heterologously expressed in *S. cerevisiae* showed that in this system, the GFP-tagged OPT3 mislocalizes to internal vesicles rather than residing at the plasma membrane (Mendoza-Cózatl et al., 2014). To prevent AtOPT3-GFP processing/folding challenges and its retention at the endoplasmic reticulum, that may occur with heterologously-overexpressed integral plasma membrane proteins that, in addition, are also fused to a relatively large epitope tag (*e.g.* GFP (Emmerstorfer et al., 2014)), we used an untagged AtOPT3 for functional complementation assays in yeast, reported in Zhai et al 2014, and this manuscript.

Here, we show that OPT3, in addition to iron and cadmium ions, also transported copper ions when expressed in *Xenopus* oocytes or the *S. cerevisiae* copper transport mutant (**Figure 3**). We also found that OPT3 was able to transport both free copper ions as well as the copper provided as Cu-NA complex (**Figure 3A**). However, it is noteworthy that OPT3-expressing oocytes accumulated 4.2 times more copper when it was provided as a free ion rather than complexed with nicotianamine, suggesting that free copper ions are the preferred OPT3 substrate in this heterologous system. Consistent with the proton-coupled nature of OPT-mediated transport, our past studies have shown that the acidification of uptake medium stimulated the OPT3-mediated inward currents (Osawa et al., 2006; Zhai et al., 2014). A recent report has shown that members of the YSL clade of the OPT transporter family, *Brachypodium distachyon* YSL3 and maize YS1 also transport copper ions in *Xenopus* oocytes (Sheng et al., 2021). Unlike AtOPT3, copper uptake by BdYSL3 was not detected when the Cu-NA complex was provided as a substrate (Sheng et al., 2021). Similar to AtOPT3, ZmYS1 was capable of transporting copper provided as free ions as well as the Cu-NA complex; similar to AtOPT3, copper ions were the preferred transport substrates of ZmYS1 in the *Xenopus* oocyte heterologous system (Sheng et al., 2021). Vascular tissue-localized NA transporters, NAET1 and NAET2, have been identified recently (Chao et al., 2021). These transporters localize to the synaptic-like vesicles in the vasculature and control iron and copper delivery to seeds. Therefore, as suggested by Chao et al 2021, OPT3 and NAET1/NAET2 may act in concert in copper/iron and NA delivery, respectively, to the phloem. In the phloem, copper and iron would then form a complex with NA to facilitate mineral delivery to the sink tissues. Similar scenario occurs with iron and citrate transport to the xylem *via* IREG1/FPN1 (**I**ron **Reg**ulated1/**F**erro**p**orti**n** 1) and FRD3 (**M**ultidrug **a**nd **T**oxin **E**fflux) transporters, respectively (Rogers and Guerinot, 2002; Green and Rogers, 2004; Morrissey et al., 2009). In the xylem, the tri-iron(III), tri-citrate complex is formed to facilitate iron delivery to shoots (Rellán-Álvarez et al., 2009).

Recent *in silico* analysis using protein secondary and tertiary structure simulation and binding affinity results of the docking analyses suggested that AtOPT3 and its counterpart from maize, ZmOPT3 may transport iron complexed with GSH (Kurt, 2021)). Despite this prediction, AtOPT3 does not transport GSH when expressed in the GSH transport mutant, *hgt1Δ,* and the addition of GSH to the *Xenopu*s oocytes bathing medium reduces cadmium transport (Zhai et al., 2014). Our finding that AtOPT3 does not transport GSH in yeast has been recently validated by (Zhang et al., 2016). AtOPT3 counterpart from rice, OsOPT7 is essential for iron homeostasis and sources to sink partitioning but does not transport Fe-NA, Fe-DMA complexes, or GSH in oocytes as well (Bashir et al., 2015). It is noteworthy that unlike AtOPT3, other OPT family members in *A. thaliana*, AtOPT4 and AtOPT6 and the closest OPT3 homolog from *Brassica juncae*, BjGt1 transport GSH (Bogs et al., 2003; Cagnac et al., 2004; Zhang et al., 2016). Recent analyses of the *Arabidopsis opt6* mutant implicated AtOPT6 in GSH long-distance transport and delivery to sink organs, especially flowers (Wongkaew et al., 2018). It is noteworthy that similar to the *opt3* mutant, the loss-of-function of At*OPT6* delays the transition from the vegetative to reproductive stage of the development (Wongkaew et al., 2018). It would be interesting to test whether mineral nutrient homeostasis is disrupted in the *opt6* mutant.

### AtOPT3 Mediates Copper (and Iron) Loading to the Phloem Companion Cells for the Delivery to Sink Tissues

The role of AtOPT3 in iron and copper loading to the phloem companion cells for subsequent partitioning from source to sink tissues is evidenced by the decreased accumulation of these metals in the phloem of the *opt3* mutant (**Figures 1F, G, K** and **Supplemental Figure S1E online** and (Zhai et al., 2014)). At the whole plant level, this function of AtOPT3 is important for the phloem-based delivery of copper and iron from sources to sinks, including roots and young leaves (**Figure 2A**, **Supplemental Figure 3 online** and (Zhai et al., 2014), respectively). Here we also show that AtOPT3 is important for copper and iron delivery to developing embryos and, in addition to leaves, silique valves might act as sources of these minerals for developing seeds (**Figure 2B** and **C**, and **Supplemental Figure 4A online**). This suggestion is based on our findings that silique valves of the mutant overaccumulated iron and copper while developing embryos accumulated less iron and copper compared to wild type (**Figure 2B** and **C**). We note that unlike copper and iron, manganese and zinc were overaccumulated in the vasculature and both silique valves and embryos, while zinc was also overaccumulated in seeds of the *opt3* mutant compared to corresponding tissues in wild type (**Figure 1H, I** and **2B**). Together, these data suggest that AtOPT3 functions in copper and iron but not manganese and zinc loading to the phloem and the phloem-based delivery to sink tissues.

We note that while the role of AtOPT3 and OsOPT7 in iron delivery to seeds is evidenced by the lower iron concentration in mature seeds (**Supplemental Figure 4B, C online** and (Stacey et al., 2008; Mendoza-Cózatl et al., 2014; Zhai et al., 2014; Bashir et al., 2015), copper accumulation in mature seeds was not affected in the *opt3* mutant in our assays using ICP-MS analysis (**Supplemental Figure 4C online** and (Stacey et al., 2008)). However, we noted a subtle difference in copper distribution in the mature seed of the *opt3* mutant *vs.* wild type using CT-XRF. Specifically, less copper was associated with the vasculature of the *opt3* mutant *vs.* wild type, and somewhat more copper was associated with the seed coat of the mutant (**Supplemental Figure 4B online**). However, this subtle difference in copper distribution was not sufficient to account for total seed concentration.

### AtOPT3 Mediates Copper Homeostasis and Systemic Copper Signaling

Our finding that OPT3 transports copper in heterologous systems and that both *opt3-2* and *opt3-3* mutant alleles accumulate less copper in roots and young leaves compared to wild type (**Figure 2A** and **Supplemental Figure 3 online**) suggested that the mutant might be more sensitive to copper deficiency and/or manifest molecular symptoms of copper deficiency. Indeed, the size of rosette leaves of both *opt3-2* and *opt3-3* mutants grown under copper deficiency was significantly smaller compared to control conditions, and their leaves were extensively chlorotic compared to wild type also grown without added copper (**Figure 4** and **F Supplemental Figure 6 online)**. Furthermore, consistent with our recent findings of the role of copper in transition to flowering (Rahmati Ishka and Vatamaniuk, 2020), the *opt3* mutant failed to flower within the time frame of the experiment and developed more rosette leaves than the wild type when grown in the medium without and even with copper (**Figure 4C, D**). Copper deficiency-associated phenotypes were rescued by transferring the mutant to a medium with higher copper concentrations (**Figure 4C** to **E** and **Supplemental Figure 7 online**).

In addition, the *opt3* mutant mounted molecular copper-deficiency responses even when it was grown under control conditions. This was evidenced by the increased activity of cupric reductase in roots and the upregulated expression of the copper-deficiency regulon in roots and young leaves of the *opt3* mutant *vs.* wild type (**Figure 4** and **Supplemental Datasets 2 to 4**). Changes in the transcriptome of mature leaves of the *opt3* mutant were indicative of neither deficiency nor sufficiency. The transcriptional copper deficiency responses of young leaves and an unspecific response of mature leaves in the *opt3* mutant are consistent with their respective copper concentrations (**Figure 2A, Supplemental Figure 3 online** and **Supplemental Datasets 2 to 4**). By contrast, copper accumulation in roots of the *opt3* mutant, although was lower than in wild type, was still at the level of sufficiency (**Figure 2A** and (Broadley et al., 2012)); nevertheless roots of the *opt3* mutant mounted the transcriptional copper deficiency response.

Using reciprocal grafting with wild type and the *opt3* mutant, we showed that OPT3 function in the shoot regulates not only iron but also copper deficiency responses of the root (**Figure 4**) suggesting the existence of shoot-to-root copper status signaling and the contribution of OPT3 to this process. Our finding that copper feeding *via* the phloem in the shoot downregulates the expression of the copper deficiency markers, *CITF1*, *COPT2* and *copper miRNAs* in the root of copper-deficient wild type and the *opt3* mutant that constitutively overexpresses copper deficiency marker genes even when it is grown under control conditions, favors the existence of OPT3-mediated systemic copper signaling (**Figure 9B** and **D**). Furthermore, our data suggest that similar to iron, phloem copper act as a repressor of copper-deficiency responsive genes in the root.

An upstream regulator of *CITF1*, *COPT2* and *copper miRNAs*, SPL7, is expressed in the vascular tissues of both shoot and roots, locally senses copper availability, and transmits it to surrounding cells (Araki et al., 2018). In the root, *SPL7* is detected in the pericycle (Long et al., 2010). Considering that phloem pole pericycle plays a major role in regulating symplastic phloem unloading in the root (Ross-Elliott et al., 2017), it is possible that SPL7 senses the availability of the phloem-arrived copper in the pericycle to regulate the expression of *CITF1*, *COPT2*, *copper miRNAs*, and, perhaps, other SPL7 targets in the root.

### Copper and Iron Can, in Part, Substitute Each Other’s Signaling Function in the Phloem

One of the unexpected outcomes from the phloem feeding experiments was finding that feeding with copper *via* the phloem in the shoot downregulated the expression of both copper and iron deficiency markers in roots of the *opt3* mutant (**Figure 9D**). Likewise, iron feeding *via* the phloem in the shoot downregulated the expression of *FIT* and *IRT1* as well as *CITF1* and *COPT2* in roots of the *opt3* mutant (**Figure 9B**). We note that the *opt3* mutant has a lower concentration of both copper and iron in the phloem and thus, provides a sensitized genetic background for the phloem feeding experiments with copper or iron. Therefore, we used wild type plants, subjected them to copper and iron deficiency individually or simultaneously, and fed with either copper or iron *via* the phloem in the shoot. We found that either copper or iron phloem feeding downregulated the expression of both copper and iron-deficiency-responsive genes (**Figure 10B and C**). These findings raised an intriguing possibility that mobile iron and copper in the phloem can, in part, substitute each other’s signaling function in regulating the expression of copper and iron uptake genes in the root (**Figure 12**).

### AtOPT3-facilitated Copper and Iron Transport to the Phloem Companion Cells Mediates Iron-Copper Crosstalk in Shoot-to-Root Signaling and Iron and Copper Homeostasis

As a corollary to finding that AtOPT3 loads copper to the phloem companion cells in the shoot was the discovery that this function is important not only for feeding the developing sinks with copper but also for conveying copper status of the shoot to the root; in addition, both copper and iron can substitute each other signaling function in the phloem of the *opt3* mutant for fine-tuning the expression of copper/iron uptake systems in the root (**Figures 9** and **12A**). Our finding that iron deficiency significantly increases the expression of already highly expressed iron uptake system in roots of the *opt3* mutant (**Figure 5D**), is especially relevant for the understanding of the OPT3-mediated copper-iron crosstalk in long distance signaling as discussed below.

We propose that in *A. thaliana* wild type grown under copper and iron sufficiency, copper and iron ions, absorbed by the root and delivered to the shoot, loaded to the phloem companion cells in leaves by AtOPT3 to convey copper/iron sufficiency (**Figure 12A**). *Under iron deficiency,* AtOPT3 loads mainly copper into the phloem companion cells since iron is limited, while copper absorption by roots and accumulation in leaves increases under iron deficiency (**Figure 12B**) and (Waters et al., 2012; Waters and Armbrust, 2013; Waters et al., 2014). While the expression of iron uptake genes in roots of iron-deficient wild type increases, it is significantly lower than in the *opt3* mutant under iron deficiency (**Figures 5D**).

Similarly, *under copper deficiency*, AtOPT3 loads mainly iron into the phloem companion cells since copper is limited (**Figure 12C**). This AtOPT3 function fine-tunes the expression of iron and copper transport systems to ensure adequate iron/copper uptake while preventing toxicity. It is noteworthy that unlike copper or iron feeding *via* the vasculature in the shoot, the function of AtOPT3 in the wild type does not entirely suppress copper or iron uptake systems in the root under reciprocal deficiency since copper or iron uptake systems are still upregulated (**Figure 12B, C**). It is possible that the concentration of iron or copper in the phloem in *A. thaliana* grown under copper or iron may not be sufficient to fully repress the expression of copper or iron uptake genes in the root. It is also possible that the sink strength and competition (roots *vs.* shoot apices and developing leaves) are also important contributing factors that determine the amount of copper and iron (as well as other components of the phloem sap) recirculated to roots *via* the phloem to repress iron or copper uptake under the reciprocal deficiency.

*Concerning the opt3 mutant*, we propose that the constitutive expression of copper and iron uptake systems in its roots originates from the constitutive copper and iron deficiency in the leaf phloem, most likely the phloem companion cells (**Figure 12D, E**). Copper and iron deficiencies, co-occurring in the phloem of the *opt3* mutant, act autonomously in regulating copper and iron deficiency genes in the roots. This suggestion is based on our finding that the expression of *CITF1* and copper *miRNAs*, normally downregulated by iron deficiency while upregulated by copper deficiency when applied independently, was upregulated under simultaneous application of both deficiencies (**Figure 10A** and **12D**). Likewise, although *FIT* is upregulated by iron deficiency and does not respond transcriptionally to copper deficiency, it was upregulated by the simultaneous application of copper and iron deficiency (**Figure 10A**).

The question remains about how copper and iron act in systemic signaling. It is possible that copper/iron availability in the phloem may be communicated *via* a common transduction pathway and/or metal-binding ligand. In this regard, iron sensors, HRZs/BTS and IDEF1 in *A. thaliana* and *O. sativa* bind not only ionic Fe^2+^ but also other transition metals and act in the root to regulate the iron uptake system (Selote et al., 2014; Kobayashi, 2019). Phloem sap contains potential iron and copper binding ligands such as NA and glutathione (GSH) (reviewed in (Gayomba et al., 2015)). It is noteworthy that the *nas4x* mutant of *A. thaliana*, lacking functional NA synthase genes, overaccumulates iron in the vasculature, similar to the *opt3* mutant (Nguyen et al., 2022). Unlike the *opt3* mutant, copper accumulation in the vasculature of the *nas4x* mutant was not observed, suggesting that in *A. thaliana* NA contributes mainly to iron homeostasis (Klatte et al., 2009; Nguyen et al., 2022). While roots of the *nas4x* mutant exhibit somewhat elevated levels of *FIT*, *IRT1* and *FRO2* expression compared to wild type both grown under control conditions, unlike the *opt3* mutant (**Figure 5D**), the expression of iron uptake genes in roots of *nas4x* does not increase further under iron deficiency (Klatte et al., 2009). It is tempting to speculate that copper loaded by OPT3 to the phloem of the *nas4x* mutant is involved in controlling the expression of iron uptake genes in its roots. The *nas4x* mutant will provide an excellent complementary system for the study of crosstalk between copper and iron in systemic signaling. A new family of phloem mobile peptides, IMA/FEP, has been proposed to coordinate systemic iron deficiency signaling. Their contribution to the crosstalk between iron and copper systemic signaling is unknown (Grillet et al., 2018; Hirayama et al., 2018).

While our data are consistent with the repressive role of the phloem-associated iron and copper in systemic signaling and the expression of iron and copper uptake systems in the root, a promotive mechanism of systemic signaling has been put forward as well (Grusak and Pezeshgi, 1996; Vert et al., 2003; Kumar et al., 2017). In this mechanism, iron-sufficient status in the shoot would prevent a release of the promotive systemic signal, and iron deficiency-responsive genes will not be expressed in the root. Considering this scenario, it is possible that iron and/or copper abundance in the phloem companion cells would sequester or inhibit synthesis of the promotive signal(s) preventing the upregulation of genes encoding the root iron and/or copper uptake systems (**Figure 12**). We note that both repressive and promotive mechanisms rely on the availability of iron and copper in the phloem, most likely the phloem companion cells.

To conclude, our studies have assigned new functions to AtOPT3 in loading copper into the phloem, subsequent distribution to sink tissues and systemic signaling of copper deficiency. We have also discovered that copper can, in part, substitute for iron in long-distance signaling. We have also shown the important role for OPT3 in copper-iron crosstalk in normal homeostasis of both elements. Understanding the basic mechanisms plants use to coordinate iron and copper demands with their uptake, transport and utilization will provide promising avenues for targeted biofortification strategies directed at increasing iron density in the edible portions of crops and improving agricultural productivity on iron and copper-deficient soils.

## Materials and Methods

### Plant Material and Growth Conditions

*Arabidopsis thaliana* (*cv*. Col-0) and a previously described *opt3-3* T-DNA insertion mutant (cv. Col-0; SALK_058794C; (Zhai et al., 2014) were obtained from ABRC. Soil-grown plants were directly sown onto Lambert 111 irrigated with regular fertilizer. For copper deficiency treatments, plants were either grown on half-strength Murashige and Skoog (MS) medium (Sigma-Aldrich, M5519) with 1% sucrose and 0.7% agar or hydroponically in a medium containing 1.25 mM KNO_3_, 0.625 mM KH2PO_4_, 0.5 mM MgSO_4_, 0.5 mM Ca(NO_3_), 10 µM Fe(III)HBED and the following micronutrients: 17.5 µM H_3_BO_3_, 3.5 µM MnCl_2_, 0.25 µM ZnSO_4_, 0.05 µM NaMoO_4_, 2.5 µM NaCl and 0.0025 µM CoCl_2_ (Arteca and Arteca 2000), and the indicated concentrations of copper with 125 nM copper considered as control (Yan et al., 2017). For growing plants hydroponically, seeds were surface sterilized before sowing onto 0.7% (w/v) agar aliquoted in 10 µL pipette tips. Pipette tips were cut before placing them into floats made of foam boards. The roots of seedlings were immersed into the hydroponic solution after 7-8 days of growth. To ensure that both wild type and the *opt3-3* mutant were treated at the same growth stage, all plants were transferred when the rosette reached to 50% of final size (principal growth stage 3.5 as documented in (Boyes et al., 2001)). In this case, wild type and the *opt3-3* plants were transferred 3 and 4 weeks after sowing seeds, respectively. The solution was replaced on a weekly basis and was replaced for one more time 24 hours before collecting samples All plants were grown in a growth chamber at 22℃, 14 h light/10 h dark photoperiod at a photon flux density of 110 µmol m^-2^ s^-1^.

### High-Throughput Sequencing of mRNA, Sequence Mapping and Differential Gene Expression Analysis

*A. thaliana* wild type and *opt3-3* plants were grown hydroponically with 0.125 μM CuSO_4_ for four weeks. The total RNA was isolated from roots, mature and young leaves using the TRIzol reagent (Invitrogen). To achieve iron deficiency, a subset of plants grown as described above were transferred to a medium without iron for four days before roots were collected. Tissues were pooled from four plants grown in the same container and three containers were used per each genotype. The strand-specific RNA-Seq libraries were constructed using 3 µg of total RNA according to procedures described previously (Zhong et al., 2011). RNA-Seq libraries were sequenced on the Illumina HiSeq 2500 system using the single-end mode. Three replicates were used for RNA-seq. Each replicate consisted of tissues (roots, young and mature leaves) pooled from four plants per genotype grown in the same container. Trimmomatic (Bolger et al. 2014) was employed to remove adaptor and low-quality sequences in the Illumina raw reads. Reads shorter than 40 bp were discarded. The resulting reads were aligned to the ribosomal RNA database using Bowtie with 3 mismatches allowed (Langmead et al., 2009; Quast et al., 2013), and those aligned were discarded. The final clean reads were aligned to the Arabidopsis genome sequence (TAIR 10) using TopHat with 1 mismatch allowed (Trapnell et al., 2009). Finally, the counts of mapped reads for each gene model from each sample were derived and normalized to RPKM (reads per kilobase of exon model per million mapped reads). DESeq2 was used to identify differentially expressed genes (DEGs) with the raw count data (Love et al., 2014). Raw *p* values were corrected for multiple testing using the false discovery rate (FDR; (Benjamini and Hochberg, 1995). Genes with an FDR less than 0.05 and fold-changes greater than or equal to 1.5 were regarded as DEGs. GO term enrichment and gene functional classification analyses were performed using Plant MetGenMap (Joung et al., 2009).

### ICP-MS Analysis of Copper Concentration in Plant Tissues

Root and shoot tissues were collected from five-week-old plants grown hydroponically as described above. The seeds and siliques were collected from 11–12-week-old plants grown in soil as described above. Iron and other elements were desorbed from root cell walls by washing roots in 10 mM EDTA for five minutes and a solution with 0.3 mM bathophenanthroline disulphonate (BPS) and 5.7 mM sodium dithionite for ten minutes followed by three consecutive washes in deionized water. The fully expanded mature leaves (three leaves/plant) and two-to-three upmost leaves less than five millimeters long (young leaves) were rinsed with deionized water three times and then pooled separately as mature leaves and young leaves. The dry seeds and valves were collected from plants grown in soil, and the seeds were separated from the valves with 500 μm mesh. Four to eight plants were pooled for one measurement in each experiment. Three measurements and three independent experiments were conducted. All the samples were dried in an 80°C oven before measuring the weight. Elemental analysis was performed using ICP-MS.

### RT-qPCR Analysis

Roots and leaves were collected from plants at Zeitgeber time 7 (Zeitgeber time is defined as the hours after lights-on) and flash-frozen in liquid nitrogen before the homogenization. Total RNA was isolated using TRIzol reagent (Invitrogen) according to the manufacturer’s instructions. cDNA templates used for qPCR analysis were synthesized by using AffinityScript QPCR cDNA synthesis kit. One μg of total RNA was treated with DNase I (New England Biolabs) prior to the first-strand cDNA synthesis to eliminate genomic DNA contamination. Real-time PCR analysis was performed in a total volume of 15 mL containing 1x iQSYBR GreenSupermix (Bio-Rad), 500 nM concentration of each PCR primer and 2 μL of 15x diluted cDNA templates using CFX96 real-time PCR system (Bio-Rad) as described (Gayomba et al., 2013). Data were normalized to the expression of *ACTIN2* (AT3G18780), whose expression was stable under investigated conditions and in studied genotypes (Supplemental Dataset 5 online). The fold difference (2^-DDCt^) and the statistical parameters were calculated using the CFX Manager Software, version 3.1 (Bio-Rad).

### Grafting Experiments

The grafting method was described previously with slight modification (Marsch-Martínez et al. 2013). All of the procedures were performed under Leica S6E microscope. Briefly, wild type and *opt3-3* seeds were germinated and grown vertically on half-strength MS medium with 0.5% sucrose for 6-7 days. Cotyledons were removed by using sterile scalpels with No.11 blades right before grafting. The scions and rootstocks were then separated and moved to half-strength MS plates with 0.5% sucrose and 0.7% agar for alignment. After grafting, the plants were grown vertically on the plates for another 10 days. Successfully grafted plants without adventitious roots above the graft junction were then transferred to the hydroponic system with 0.25 μM CuSO_4_ for 24 days before qPCR analysis.

### Synchrotron X-Ray Fluorescence Imaging

Fresh samples were detached immediately before the analysis and were placed in a wet chamber made between two layers of metal-free Kaptonfilm and mounted onto 35-mm slide frames. The spatial distribution of Cu and Fe was imaged via SXRF microscopy at the F3 station at the Cornell High Energy Synchrotron Source (CHESS). The 2D Cu and Fe raster maps were acquired at the 25-µm resolution, 0.2 s/pixel dwell time using a focused, monochromatic incident x-ray beam at 12.8 keV and photon flux of ∼10^10^photons/s. The monochromatic beam was generated with 0.6% energy bandwidth multilayers. Focusing was achieved using a single-bounce moncapillary (named PEB605) fabricated at CHESS. These settings did not cause detectable damage to plant tissues within the 6- to 9-h scans required for analysis of the full set of samples. Element-specific x-ray fluorescence was detected using a Vortex ME-4 silicon drift detector. Quantifications were done by calibrating using a thin copper foil film standard (15.9 µg copper/cm^2^, the metal was deposited on 6.3 µm thick Mylar) during each experiment and concentrations were expressed as µg cm^-2^. Data was processed with the software Praxes, which was developed at CHESS and employs PyMCA libraries in batch mode (Solé et al., 2007).

### Synchrotron-based Confocal XRF Microscopy

Confocal XRF experiments were obtained at beamline 5-ID (SRX). The beamline monochromator and focusing optics were employed to deliver 3×10^11^ photons/second in a 1×1 µm^2^ beam, with incident beam energy of 10.0 keV and bandwidth of approximately 1 eV. The confocal geometry was achieved by placing a collimating channel array (CCA) (Agyeman-Budu et al., 2016) onto a single-element Vortex SDD detector perpendicular to the beam. The sample stage was oriented such that the horizontal translation axis of the stage is 35 degrees from that of the incident beam, and 55 degrees from the detector axis. The particular CCA employed for these measurements has a series of 175, 2-µm channels etched into a 1-mm thick Germanium substrate, etched to an approximate depth of 300 µm. The working distance between the sample and optic is 1.5 mm. Due to the finite width of the optic at its front tip, the maximum probe depth of the system in this configuration is 0.8 mm. Quantitative calibration of the confocal XRF system was achieved by methods described in (Malzer and Kanngießer, 2005; Mantouvalou et al., 2012).

### Synchrotron-based X-ray Fluorescence Computed Microtomography (F-CMT)

Internal distributions of copper, iron and other micronutrient elements in wild type and the *opt3-3* mutant seeds were measured *in vivo* by synchrotron-based X-ray Fluorescence Computed Microtomography (F-CMT) at the X-ray Fluorescence Microprobe (XFM) beamline at the National Synchrotron Light Source II (NSLS-II) in Upton, NY. XFM (4-BM) beamline is designed for monochromatic operation in the 2.3 to 23 keV range and optimized for high-quality, spatially-resolved X-ray absorption spectroscopy (Sulfur to Technetium K-edges) in conjunction with element-specific imaging and microdiffraction. XFM beamline can also be operated in a pink beam “imaging” mode that delivers a 1-micron spot with up to 1000X more flux than the XFM monochromatic beam. XFM filtered pink beam (12 – 20 keV broadband) was focused by Kirkpatrick-Baez (KB) mirrors to a 1-micron spot for F-CMT measurements of seeds. Seeds were mounted to a quartz post that interfaces with a Huber goniometer head on a rotation stage attached to a fast-scanning translation stage. F-CMT images were acquired at the seed center by rotating and translating the specimen in the microbeam while recording the fluorescence intensities with a Hitachi 4-element Vortex SDD coupled to Quantum Detectors Xspress3 electronics. F-CMT data were collected using 0.75 – 3.0^-^ degree angular steps, 2 – 8 µm translation steps and 50 ms dwell time. Tomographic image reconstructions using a conventional filtered back projection algorithm were processed using tomo-Py plugin to GSE Mapviewer in the LARCH package (Newville, 2013). Thin-film standard reference materials (SRM 1832 & 1833) were measured as part of the data set to establish elemental sensitivities (counts per second per mg cm^-2^). The intensity of elements between wild type and the *opt3* mutant was set to the same scale.

### Copper Uptake in *Xenopus laevis* Oocytes

Oocyte harvesting, cRNA synthesis, microinjections, and OPT3 expression in *Xenopus* oocytes was performed as described previously (Zhai et al., 2014). All animal procedures were performed in accordance with Cornell University IACUC Protocol number 2017-0139. *X. laevis* oocytes were injected with 50 nL of water (control) or 50 nL of water containing 50 ng of OPT3 cRNA and incubated in ND96 solution at 18°C for 4 d prior to the uptake measurements. The basal uptake solution consisted of a modified ND96 solution containing 96 mM NaCl, 1 mM KCl, 0.9 mM CaCl_2_, buffered with 5 mM 2-(N-morpholino) ethanesulfonic acid/NaOH to pH 6.0, as previous studies determined these conditions were suitable to minimize endogenous transport in oocytes (Zhai et al., 2014; Sheng et al., 2021). The uptake solution was supplemented with 25 µM Cu-NA or 100 µM CuSO4 [the final free Cu^2+^ activity in the uptake solution being estimated to be 35 µM as determined by GEOCHEM-EZ (Shaff et al., 2010). Each sample contained 8 – 10 oocytes, with 5 replicates per data point. At a given time point, the uptake was terminated by washing oocytes six consecutive times with an ice-cold basal uptake solution. Oocytes were collected and samples were digested in 100 µL of 70% HClO4, re-suspended in 5 ml of 0.1 M nitric acid, and analyzed using inductively coupled plasma mass spectrometry (Sciex ICP-MS). Resting membrane potentials were measured as described previously(Zhai et al., 2014).

### Functional Complementation Assays in *Saccharomyces cerevisiae*

S. cerevisiae SEY6210 (MATa ura3-52 leu2-3,-112 his3Δ200 trp1Δ901 lys2-801 suc2Δ9) wild type and the ctr1Δctr2Δctr3Δ triple mutant (MATa ura3-52 his3Δ200 trp1-901 ctr1::ura3::Knr ctr2::HIS3 ctr3::TRP1) were used for functional complementation assays. YES3-Gate-OPT3 construct or YES3-Gate lacking the cDNA insert were transformed into appropriate yeast line using the Frozen-EZ Yeast Transformation II kit (Zymo Research). Transformants were selected for uracil prototrophy on YNB medium containing 6.7% (w/v) yeast nitrogen base without amino acids (Difco), 0.77% (w/v) CSM-URA, 0.5% (w/v) NaCl, 2% glucose, 2% (w/v) agar.

Functional complementation assays included analyses of the respiration competence and copper accumulation in yeast cells as described (Jung et al., 2012). Specifically, the respiration competence of different cell lines was tested by the ability of cells to grow on non-fermentable carbon sources (Dancis et al., 1994). To do so, different cell lines transformed with *YES3-Gate* with or without *OPT3* cDNA insert were grown in liquid YNB-URA to an OD_600 nm_ = 1.0, serially 10-fold diluted, and spotted onto YPEG medium containing 1% (w/v) yeast extract, 2% (w/v) bacto-peptone, 3% (v/v) glycerol, 2% (v/v) ethanol, and 2% (w/v) agar and the indicated concentrations of CuSO_4_. Plates were incubated for 3 days at 30°C.

Different yeast lines were grown to exponential log phase in liquid YNB media described above to analyze copper accumulation. An aliquot (150 µl) of the overnight grown culture was inoculated into 20 ml of the fresh YNB media with 20 µM CuSO_4_ and cells were grown at 30°C. After 24 h, cells were collected by centrifugation, washed with deionized water before copper was desorbed from yeast cell wall by washings in desorbing buffer containing 1mM EDTA and 100 µM of a copper chelator, bathocuproine disulfonate BCS, pH8.0. Cells were then washed two more times in de-ionized water and collected by centrifugation. The cell pellet was dried, digested by heating with a combination of purified concentrated nitric and perchloric acids, and finally dissolved in 10 ml of 5% nitric acid. The concentration of copper in processed yeast cells was analyzed by inductively coupled plasma mass spectroscopy (ICP-MS; Agilent 7500).

### Collection of the Phloem Sap

Phloem sap was collected from wild type and *opt3-3* mutant plants grown hydroponically at the late vegetative stage as described (Zhai et al., 2014). Briefly, leaves # 9 and 10 collected from one plant were pooled together, and the xylem sap was flashed by placing the petioles in a tube filled with 300 µL of deionized water and incubated in an illuminated growth chamber for 15 min followed by further incubation in darkness for 1 h. To collect phloem sap, the petioles were then recut in 5 mM Na_2_-EDTA (pH 7.5) under low light before placing the petioles in pre-weighed Eppendorf tubes containing 250 µL of 5 mM Na_2_-EDTA (pH 7.5). Leaves were then incubated in darkness for 1 h in a high-humidity chamber lined with wet paper towels and sealed with Vaseline. Samples with the collected phloem sap were weighed again to determine the volume of phloem sap obtained per sample to calculate concentrations of copper or potassium (abundant in the phloem) after ICP-MS.

### Copper and Iron Feeding to the Phloem

The phloem feeding procedure was performed as described by (Oparka et al., 1994; Oparka et al., 1995; Knox, 2019). The procedure is based on the abrasion of the source leaf vein that results in some phloem ‘trapping’ of the applied sample. To test the phloem loading effectiveness, we used a phloem tracer carboxyfluorescein diacetate succinimidyl ester (CFDA), that after diffusing into plant cells, is cleaved by cellular esterases into a membrane-impermeable form, carboxyfluorescein (CF) (Grignon et al., 1989). *A. thaliana* wild type was grown hydroponically in control conditions for 33 days. Ten microliters of 10 mM CFDA was applied to the leaf surface around the major vein of six to seven bottommost leaves before the leaf surface at the major vein was abrased with a carbon steel blade. CF-mediated fluorescence in roots was visualized after 6 hours using a FITC filter set of the Axio Imager M2 microscope equipped with the motorized Z-drive (Zeiss). Images were collected using the high-resolution 25 AxioCam MR Camera and processed using the Adobe Photoshop software package, version 12.0.

For iron or copper phloem feeding, *A. thaliana* wild type was grown hydroponically with or without 125 nM CuSO_4_ for 33 days. To impose iron deficiency, plants were grown hydroponically in a control solution for 31 days and then a subset was transferred to a fresh medium without iron and grown for two more days. The *opt3* mutant was grown hydroponically under control conditions for 33 days. Ten microliters of 5 mM CuSO_4_ or 10 mM FeCl_2_ dissolved in 10 mM citrate were applied to the major vein of 6 to 7 bottommost leaves before the leaf surface at the major vein was abrased with a carbon steel blade. For control, water (a mock treatment for CuSO_4_) or 10 mM citrate (a mock treatment for Fe(II)-citrate) were fed to the phloem as described above. This procedure was performed on 8 fully developed leaves per each plant. In all experiments, roots were collected after 24 hours of phloem feeding and at Zeitgeber time 4-5. Roots were pooled from four plants per experiment and all experiments were repeated at least three times.

## Statistical analysis

Statistical analyses of experimental data were performed using the ANOVA single-factor analysis and Tukey HSD using JMP^®^ Pro 14 (SAS Institute Inc., Cary, NC, 1989-2007).

## Accession numbers

Sequence data of the genes from this article can be found in the Arabidopsis Genome Initiative or GenBank/EMBL databases under the following accession numbers: *OPT3* (AT4G16370), *CITF1* (AT1G71200), *FRO4* (AT5G23980), FRO5 (AT5G23990), *COPT1* (AT5G59030), *COPT2* (AT3G46900), *YSL1* (AT4G24120), *YSL2* (AT5G24380), *YSL3* (AT5G53550), *CSD1* (AT1G08830), *CSD2* (AT2G28190), *IRT1* (AT4G19690), *FRO2* (AT1G01580), *FRO6* (AT5G49730), *FRO7* (AT5G49740), *FSD1* (AT4G25100), *ARPN* (AT2G02850), *BCB* (AT5G20230), *CCS1* (AT1G12520), *UCC2* (AT2G44790), *FER1* (AT5G01600), *FER3* (AT3G65090), *FER4* (AT2G40300), *ZIP2* (AT5G59520), *bHLH23* (AT4G28790), TCH4 (AT5G57560), *BTSL1* (AT1G74770), *ZAT12* (AT5G59820), *FEP2/IMA2* (AT1G47395), *FEP1/IMA3* (AT2G30766), *IREG2/FPN2* (AT5G03570), *VTL1* (AT1G21140), *VTL2* (AT1G76800), *VTL5* (AT3G25190), *NAS3* (AT1G09240), *PETE2* (AT1G20340), *MT1B* (AT5G56795), *NRT2.7* (AT5G14570), *MT1B* (AT5G56795), *MT1A* (AT1G07600), *MT2A* (AT3G09390), *AT2G47010, AT4G10500, At1G31710, At1G32350, At5G02670,* and *At5g05250*.

## Author Contributions and Acknowledgements

J-C. C. and O. K. V. designed experiments; all authors contributed to different experiments presented in this manuscript. J.C. and O. K. V. wrote the manuscript with contributions from all authors. We thank Mary Lou Guerinot (Dartmouth, USA) for providing the seed of the *fit*-*2* mutant and for constructive comments on the manuscript. We thank David Mendoza-Cózatl (University of Missouri) for providing seeds of the *opt3-2* mutant. We thank a former member of the Vatamaniuk lab, Nanditha Vimalakumari for her contribution to Figure 3 and all current lab members for constructive comments on the manuscript. We thank Prof. Margaret Frank and her Ph.D. student, Hannah Thomas (Cornell University) for assisting with the phloem feeding experiments. We thank Dr. John Grazul at the Cornell Center for Materials Research (CCMR) for assisting in the preparation samples for 2D-CXRF. The CCMR facility is supported by the National Science Foundation under Award Number DMR-1719875. Parts of this research used the XFM and SRX Beamlines of the National Synchrotron Light Source II, a U.S. Department of Energy (DOE) Office of Science User Facility operated for the DOE Office of Science by Brookhaven National Laboratory under Contract No. DE-SC0012704. This work is based upon research conducted at the Cornell High Energy Synchrotron Source (CHESS), which during the period of research was supported by the National Science Foundation under award DMR-1332208; the Center for High Energy X-ray Sciences (CHEXS) is presently supported by the National Science Foundation under award DMR-1829070. This study was funded by NSF-IOS #1656321 to O. K. V. and NSF-IOS #1754966 to E. W., O. K. V. and M. P.

---

**Supplemental Figure S1 (Supports Figs. 1 and 2). Copper and iron maps and copper accumulation in the phloem exudates of wild type and the *opt3-3* mutant.**

(**A**) and (**B**) show the representative leaves used for 2D-XRF analysis. Plants were grown hydroponically with 250 nM CuSO_4_ for 22 days before 2D-XRF analysis. Numbers indicate the order of leaf from the oldest to the youngest. Red arrows indicate the fourth oldest true leaves that were analyzed in Figure 1. (**C**) and (**D**) show the 2D-XRF mapping of the 1^st^ true leaf (**C**) and the 6^th^ true leaf (**D**) of the wild type (***Wt***) and the *opt3* mutant (***opt3***) grown hydroponically with 250 nM Cu for 24-26 days. Bars = 1 mm. (**E**) ICP-MS analysis of the phloem sap collected from the wild-type and the *opt3* mutant. A minimum of ten individual phloem sap collections per genotype were performed. Plants were grown and phloem sap was collected as described in (Kumar et al., 2017).

**Supplemental Figure S2 (Supports Fig. 2). The *opt3-3* mutant accumulates a high concentration of metals in vegetative tissues.**

Plants were grown hydroponically with 125 nM (**A**) or 250 nM CuSO_4_ (**B**) for 30 days before tissue collection for ICP-MS analysis. Shown values are arithmetic means ± S.E. Asterisks indicated statistically significant differences from wild type (*p* < 0.05, Student’s *t* test, n = 3 independent experimental set-ups. In each experimental set-up, tissues from four to five plants grown in the same container were pooled and represented an independent measurement.

**Supplemental Figure S3 (Supports Fig. 2). The *opt3-2* mutant accumulates less copper in roots and young leaves.**

Plants were grown hydroponically with 125 nM CuSO_4_ (**B**) for 30 days before tissue collection for ICP-MS analysis. Shown values are arithmetic means ± S.E. Asterisks indicated statistically significant differences from wild type (*p* < 0.05, Student’s *t* test, n = 3 independent measurements. In each measurement, tissues were pooled from four to five plants grown in the same container).

**Supplemental Figure S4 (Supports Fig. 2). OPT3 mediates copper and iron accumulation in developing embryos.**

**(A)** Fifteen-mm-long developing siliques were collected from soil-grown plants and subjected to 2D-SXRF analysis. (**B**) Dry seeds collected from soil-grown wild-type and the *opt3-3* mutant were subjected to 2D CT-XRF. Blue arrows point to the vasculature, blue boxes to regions in the seed coat with contrasting copper accumulation. A representative image of at least three scanned specimens is shown. (**C**) ICP-MS analysis in seeds of wild type and the *opt3-3* mutant. Shown values are arithmetic means ± S.E. Asterisks indicate statistically significant differences from wild type (*p* < 0.05, Student’s *t* test, n = 3 independent experimental set-ups. In each set-up, tissues were pooled from four to five plants to yield one ICP-MS measurement).

**Supplemental Figure S5 (Supports Fig 3A). Cellular integrity of mock and OPT3-expressing oocytes in control or copper-containing solutions.**

Cells were (**A**) exposed to a bath media containing 100 µM CuSO_4_ for 3 hours and subsequently washed and allowed to recover in a bath media lacking copper for the remaining 21 hours, or (**B**) continuously exposed to a bath media containing 50 µM CuSO_4_ during 24 hours. Cell integrity was monitored by measuring the cell’s membrane potentials (bottom panel) and changes in morphological characteristics (top panel) over 24 hours at the end of the 3, 9, and 24 h time intervals denoted above each panel. Copper-induced cellular damage was visually evidenced by an early blotchiness of the dark pigment in the animal hemisphere and eventual complete discoloration of the cell, predominantly in OPT3 expressing cells. Survival rates (middle row pane) were calculated as the % of damaged-looking cells at a given time point. Resting membrane potentials were measured in at least 4 different cells for each group at each given time point and were recorded as described previously (Zhai et al., 2014). Data for mock and OPT3-expressing cells exposed to copper treatment are color-coded blue and red, respectively. Mock and OPT3-expressing cells maintained in bath media lacking copper for the entire 24 hours of the experiment (labelled as non-treated) are color-coded black and grey, respectively.

**Supplemental Figure S6 (Supports Fig. 4). Seedlings of the *opt3* mutant are sensitive to copper deficiency.**

(**A**) A representative image of increased sensitivity of the *opt3-2* mutant allele to copper deficiency. Plants were grown hydroponically with or without copper supplements for 35 days. (B) to (**E**) The *opt3-3* mutant and wild type were grown on ½ MS solid medium with or without 250 µM or 500 µM of a copper chelator BCS. (**B**) shows a representative image of plants after ten days of growth, after which root length (**C**), fresh weight (**D**) and cupric reductase activity (**E**) were analyzed. In **C** to **E** data are means ± S.E. Levels not connected by the same letter are significantly different (*p* < 0.05, Tukey-Kramer HSD test). In **C**, three independent experiments were performed with 30 plants per genotype analyzed per experiment (n = 90); in **D,** n = 3 independent experiments with 30 to 40 seedlings pooled for analyses in each experiment; in **E**, n = 3 independent experiments with 20 seedlings analyzed in each experiment.

**Supplemental Figure S7 (Supports Fig. 4). Transferring the *opt3-3* mutant to high copper rescues its growth.**

(**A**) Wild type and the *opt3* mutant were germinated and grown hydroponically at 125 nM Cu for four weeks before transferring to a fresh hydroponic medium with 500 nM (**A**) or 250 nM (**B**) copper. Images were taken after one week of growth. A representative set-up/result from three independent experiments is shown.

**Supplemental Figure S8 (Supports Fig. 5). The *opt3* mutant accumulates IRT1 protein in roots under Fe sufficient growth conditions.**

(**A**) Western blots analysis of IRT1 protein accumulation in roots of wild type and the *opt3-3* mutant. Plants were grown hydroponically with 125 nM CuSO_4_ and 10 mM Fe-HBED for four weeks before tissue collection. (**B**) Western blots analysis of iron deficiency-induced IRT1 accumulation in wild type roots. Plants were grown hydroponically with 125 nM CuSO_4_ and 10 µM Fe-HBED for three weeks and then transferred to a fresh medium without iron. Roots were collected after one week of growth under iron deficiency. The *fit-2* mutant was incorporated as a negative control for the IRT1 protein accumulation. In (**A**) and (**B**), the IRT1 signals are recognized by the antibody at 35 kD while actin (protein loading control) at 45 kD. For the immunodetection of actin epitope, the nitrocellulose blots were probed with the primary mouse-monoclonal anti-actin antibody (1:5,000 dilution, Sigma-Aldrich) and secondary, an HRP-conjugated goat-antimouse IgG antibody (1:10,000 Rockland Immunochemicals). In both cases, immunoreactive bands were visualized with Clarity Max ECL blotting substrates (BIORAD).

**Supplemental Figure 9 (Supports Fig. 9). Phloem feeding with carboxyfluorescein diacetate succinimidyl ester (CFDA) in leaves of wild type *Arabidopsis thaliana*.**

(**A**) The illustration of phloem feeding procedures as described in Materials and Methods. (**B)** Shows a representative image of carboxyfluorescein (**CF**)-mediated fluorescence and brightfield images of the root after feeding 10 mM CFDA *via* the phloem in the shoot. (**C**) Shows autofluorescence of mock-fed plants (**Buffer**). (**D**) shows that iron but not mock feeding via the phloem in the shoot of iron-deficient wild type eliminates degreening of young leaves (indicated by white arrows). In all cases, experiments were repeated at least three times with at least three planst analyzed per each experiment.

**Supplemental Figure 10 (Supports Fig. 11). Iron availability in the medium influence the expression of copper deficiency responsive genes in the roots of *A. thaliana*.**

(A) Shows a representative phenotype of *A*. *thaliana* grown hydroponically with 10 µM or 50 µM FeHBED for five weeks. Twelve plants per condition were analysed in total in three independent experiments. (**B**) The accumulation of iron in roots and shoots of plants grown as in (**A**). Mean values ± S.E (n = 5 ICP-MS measurements obtained from two batches of independently grown plants per condition. Tissues from four plants were pooled for each measurement). Asterisks indicate statistically significant differences (*p* < 0.05, Student-*t* test).

