## Supplemental Information Online for "OPT3 Transports Copper to the Phloem, Mediates Shoot-to-Root Copper Signaling and Crosstalk Between Copper and Iron Homeostasis in *A. thaliana*"


**Short title:** AtOPT3 mediates systemic copper and iron signaling

**One-sentence summary:** AtOPT3 loads copper and iron into the phloem companion cells, for subsequent distribution to sink tissues and systemic signaling of copper and iron deficiency.

The author responsible for distribution of material integral to the findings presented in this article in accordance with the policy described in the Instructions for Authors (www.plantcell.org) is Olena Vatamaniuk

**Supplemental Table 1.** A list of used oligos for Qrt-PCR

| **Oligo name** | **Gene** | **Primer sequence (5' to 3')** |
|---|---|---|
| Actin-F | *Actin* | GACCTTTAACTCTCCCGCTA |
| Actin-R | *Actin* | GGAAGAGAGAAACCCTCGTA |
| CITF1-F | *CIT1* | ACGAGGTCCTTCTATTCGAGCA |
| CITF1-R | *CIT1* | ACCCTTTGCTCTCGGCAAACTT |
| COPT1-F | *COPT1* | CATGTCGTTTAACGCCGGTGTGTT |
| COPT1-R | *COPT1* | CCGGAAAGTTTGGCTTCCGAACAA |
| COPT2-F | *COPT2* | TGGTGATGCTCGCTGTTATGTCCT |
| COPT2-R | *COPT2* | TCTGGTCATCGGAGGGTTTCTTGA |
| FRO4-F | *FRO4* | TTCCAGTGTAGTTTTCTTA |
| FRO4-R | *FRO4* | TTGTACCTGATTCTTGAAC |
| FRO5-F | *FRO5* | TACCCAAAAGCAACCCTCCC |
| FRO5-R | *FRO5* | ATCATCACCCTCCCCATTCT |
| FSD1-R | *FSD1* | ACTTACAGCTTCCCAAGACAC |
| FSD1-F | *FSD1* | TGCTGTGAATCCCCTTGTG |
| CSD1-R | *CSD1* | TTCTGGCCTTAAGCCTGGTC |
| CSD1-F | *CSD1* | CGACATGCTGGTGATCTAGG |
| CSD2-R | *CSD2* | CATGACACACGGAGCTCCAG |
| CSD2-F | *CSD2* | GAGCTGGAGGGCTATATCCG |
| IRT1-F | *IRT1* | ACCCGTGCGTCAACAAAGCTAAAG |
| IRT1-R | *IRT1* | TCCCGGAGGCGAAACACTTAATGA |
| FRO2-F | *FRO2* | TGTGGCTCTTCTTCTCTGGTGCTT |
| FRO2-R | *FRO2* | TGCCACAAAGATTCGTCATGTGCG |

### Materials and Methods

#### Protein extraction and Western blot analysis

The procedures of total protein extraction were modified from Sivitz, et al., 2011. One-hundred mg of fresh root samples were frozen in liquid nitrogen and ground with a bead mill homogenizer (Omni Bead Ruptor 12). The total proteins were then extracted from the ground tissues by adding 300 μL protein extraction buffer containing 5% SDS, 5% β-mercaptoethanol, 50 mM Tris HCl (pH 7.4), 1x protease inhibitor cocktail (Sigma-Aldrich, P9599), 40 μM MG-132, (Sigma-Aldrich), and 2 mM phenylmethylsulfonyl fluoride (PMSF, Millipore-Sigma). The protein extracts were centrifuged at 13,000 rpm for 15 min, and the total soluble proteins were separated from cell debris. The samples were then boiled at 95ºC for 10 min before electrophoresis. For each sample, 5 μL total proteins were separated on a 12% sodium dodecyl sulfate polyacrylamide gel electrophoresis (SDS-PAGE) and were transferred onto a nitrocellulose membrane (BIORAD) by electroblotting. The Western blotting procedures were described previously with slight modification (Kim et al. 2010). The membranes were first blocked in EveryBlot Blocking Buffer (BIORAD). For immunodetection of IRT1 protein, the nitrocellulose blots were probed with the primary goat polyclonal anti-IRT1 antibody (1:5,000 dilution, Agrisera AS11-1780), and with the secondary HRP-conjugated antigoat-IgG antibody (1:10,000, Rockland Immunochemicals). For the immunodetection of actin epitope, the nitrocellulose blots were probed with the primary mouse-monoclonal anti-actin antibody (1:5,000 dilution, Sigma-Adrich) and secondary, an HRP-conjugated goat-antimouse IgG antibody (1:10,000 Rockland Immunochemicals). In both cases, immunoreactive bands were visualized with Clarity Max ECL blotting substrates (BIORAD).

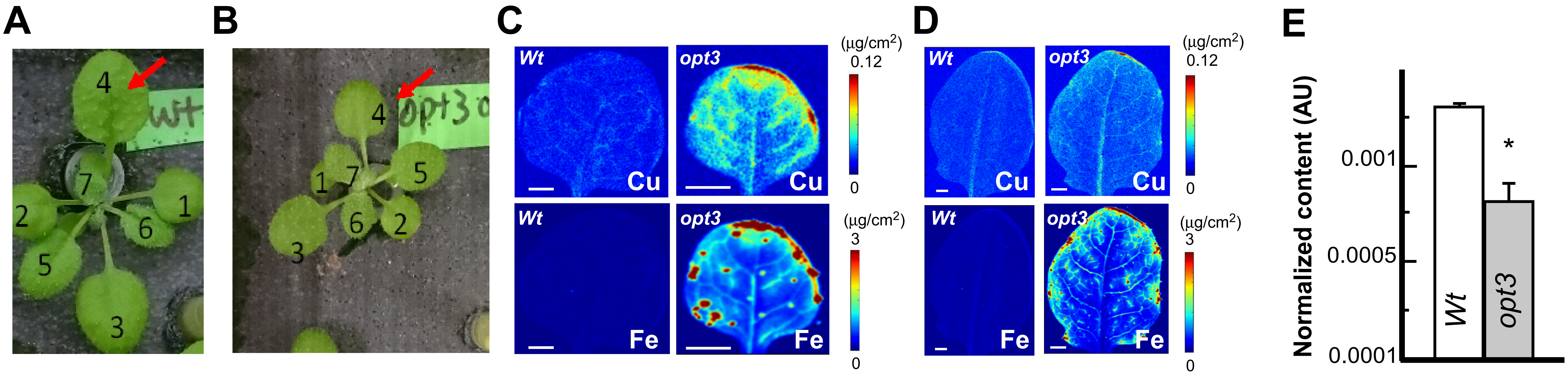


**Supplemental Figure S1 (Supports Figs. 1 and 2). Copper and iron maps and copper accumulation in the phloem exudates of wild type and the *opt3-3* mutant.**

(**A**) and (**B**) show the representative leaves used for 2D-XRF analysis. Plants were grown hydroponically with 250 nM $CuSO_4$ for 22 days before 2D-XRF analysis. Numbers indicate the order of leaf from the oldest to the youngest. Red arrows indicate the fourth oldest true leaves that were analyzed in Figure 1. (**C**) and (**D**) show the 2D-XRF mapping of the $1^{st}$ true leaf (**C**) and the $6^{th}$ true leaf (**D**) of the wild type (**Wt**) and the *opt3* mutant (***opt3***) grown hydroponically with 250 nM Cu for 24-26 days. Bars = 1 mm. (**E**) ICP-MS analysis of the phloem sap collected from the wild-type and the *opt3* mutant. A minimum of ten individual phloem sap collections per genotype were performed. Plants were grown and phloem sap was collected as described in (Kumar et al., 2017).

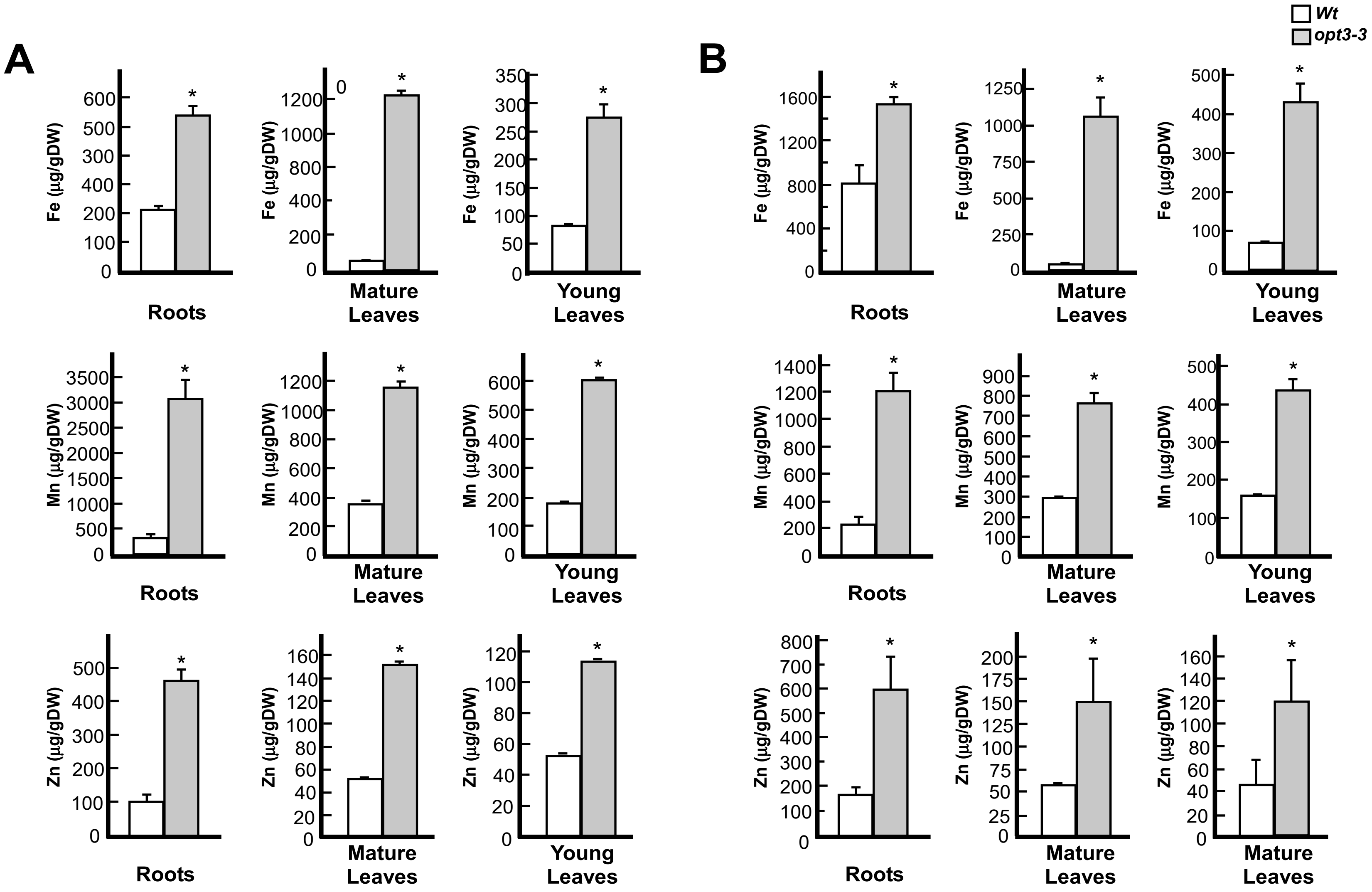


**Supplemental Figure S2 (Supports Fig. 2). The *opt3-3* mutant accumulates a high concentration of metals in vegetative tissue**s.

Plants were grown hydroponically with 125 nM (**A**) or 250 nM $CuSO_4$ (**B**) for 30 days before tissue collection for ICP-MS analysis. Shown values are arithmetic means ± S.E. Asterisks indicated statistically significant differences from wild type ($p < 0.05$, Student's $t$ test, n = 3 independent experimental set-ups. In each experimental set-up, tissues from four to five plants grown in the same container were pooled and represented an independent measurement.

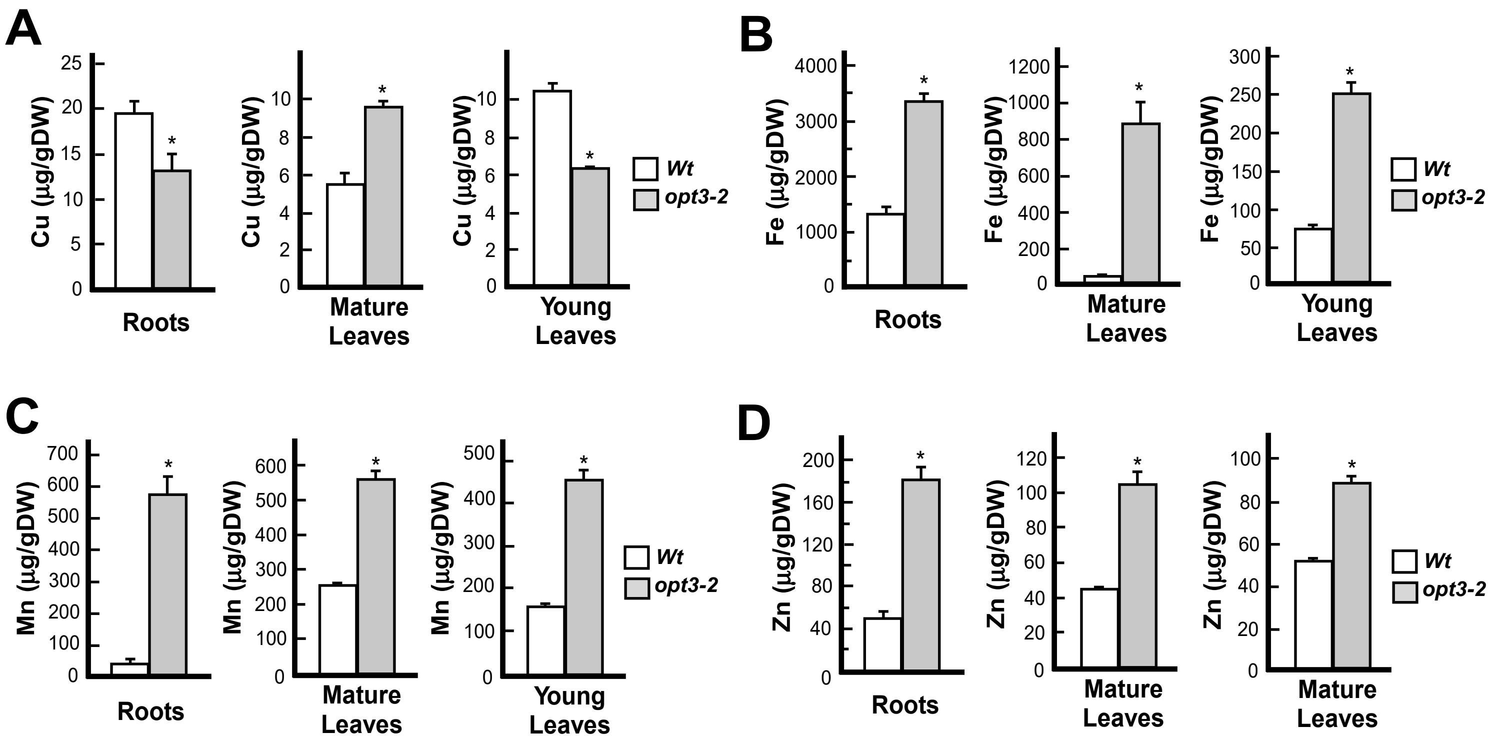


**Supplemental Figure S3 (Supports Fig. 2). The *opt3-2* mutant accumulates less copper in roots and young leaves.** Plants were grown hydroponically with 125 nM $CuSO_4$ (**B**) for 30 days before tissue collection for ICP-MS analysis. Shown values are arithmetic means ± S.E. Asterisks indicated statistically significant differences from wild type ($p < 0.05$, Student's *t* test, n = 3 independent measurements. In each measurement, tissues were pooled from four to five plants grown in the same container).

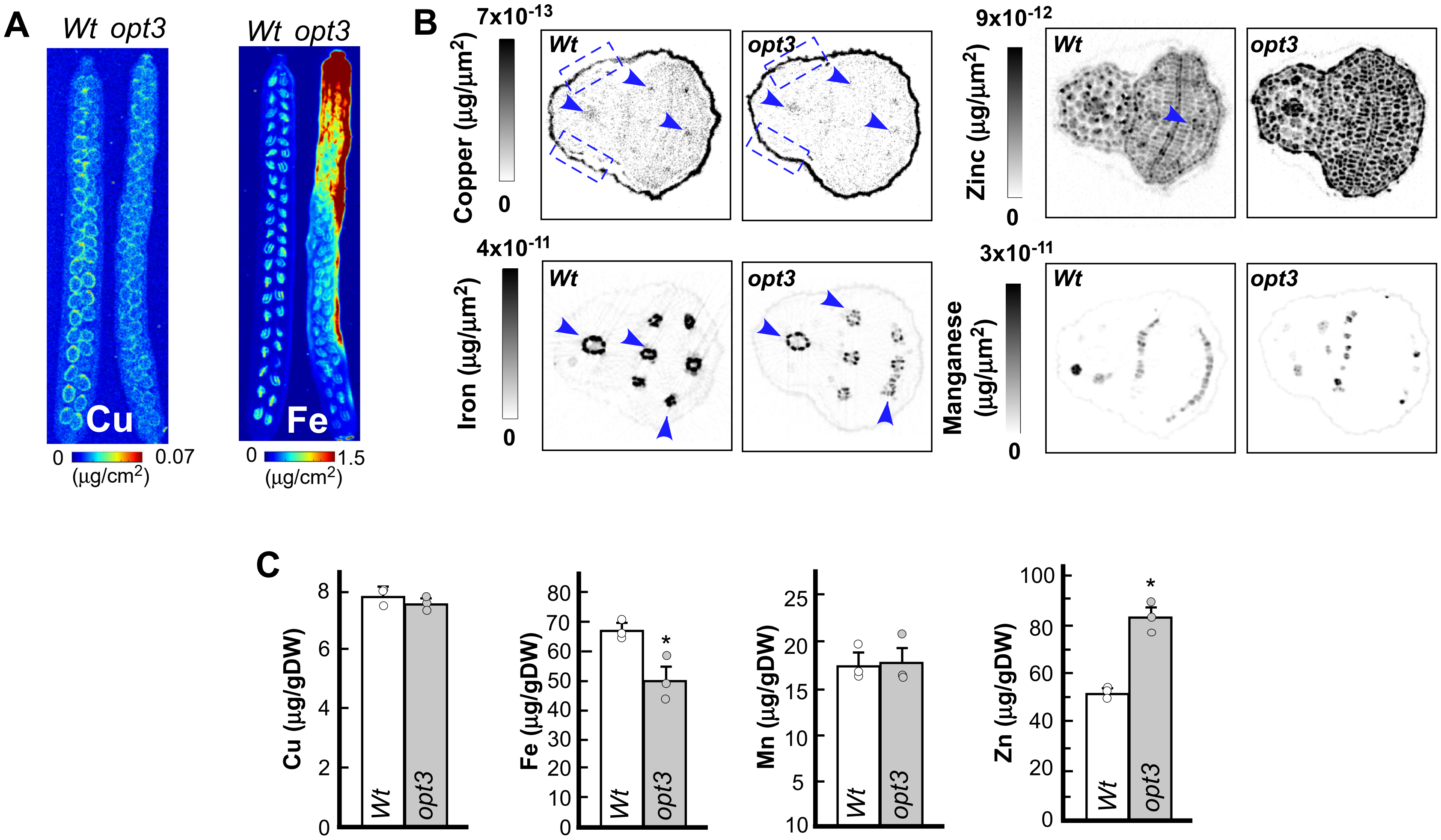


**Supplemental Figure S4 (Supports Fig. 2). OPT3 mediates copper and iron accumulation in developing embryos.**

**(A)** Fifteen-mm-long developing siliques were collected from soil-grown plants and subjected to 2D-SXRF analysis. (**B**) Dry seeds collected from soil-grown wild-type and the *opt3-3* mutant were subjected to 2D CT-XRF. Blue arrows point to the vasculature, blue boxes to regions in the seed coat with contrasting copper accumulation. A representative image of at least three scanned specimens is shown. (**C**) ICP-MS analysis in seeds of wild type and the *opt3-3* mutant. Shown values are arithmetic means ± S.E. Asterisks indicate statistically significant differences from wild type ($p < 0.05$, Student's $t$ test, n = 3 independent experimental set-ups. In each set-up, tissues were pooled from four to five plants to yield one ICP-MS measurement).

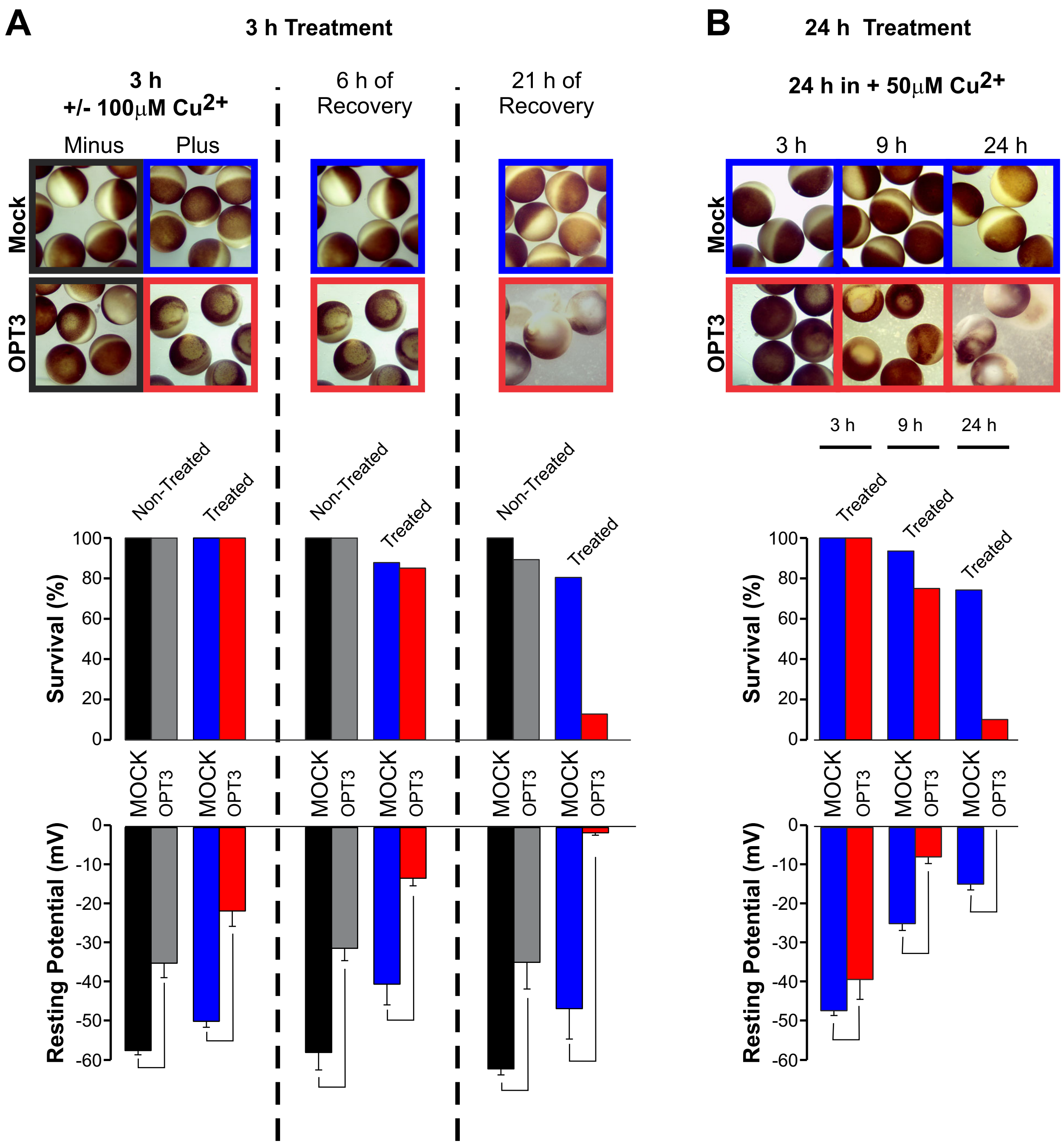


**Supplemental Figure S5 (Supports Fig. 3A) . Cellular integrity of mock and OPT3 expressing oocytes in control or copper-containing solutions.** Cells were (**A**) exposed to a bath media containing 100 µM $CuSO_4$ for 3 hours and subsequently washed and allowed to recover in a bath media lacking copper for the remaining 21 hours, or (**B**) continuously exposed to a bath media containing 50 µM $CuSO_4$ during 24 hours. Cell integrity was monitored by measuring the cell's membrane potentials (bottom panel) and changes in morphological characteristics (top panel) over 24 hours at the end of the 3, 9 and 24 h time intervals denoted above each panel. Copper-induced cellular damage was visually evidenced by an early blotchiness of the dark pigment in the animal hemisphere and eventual full discoloration of the cell, predominantly in OPT3 expressing cells. Survival rates (middle row pane) were calculated as the % of damaged-looking cells at a given time point. Resting membrane potentials were measured in at least 4 different cells for each group at each given time point and were recorded as described previously (Zhai et al., 2014). Data for mock and OPT3-expressing cells exposed to copper treatment are color-coded blue and red, respectively. Mock and OPT3-expressing cells maintained in bath media lacking copper for the entire 24 hours of the experiment (labeled as Non-treated) are color-coded black and grey, respectively.

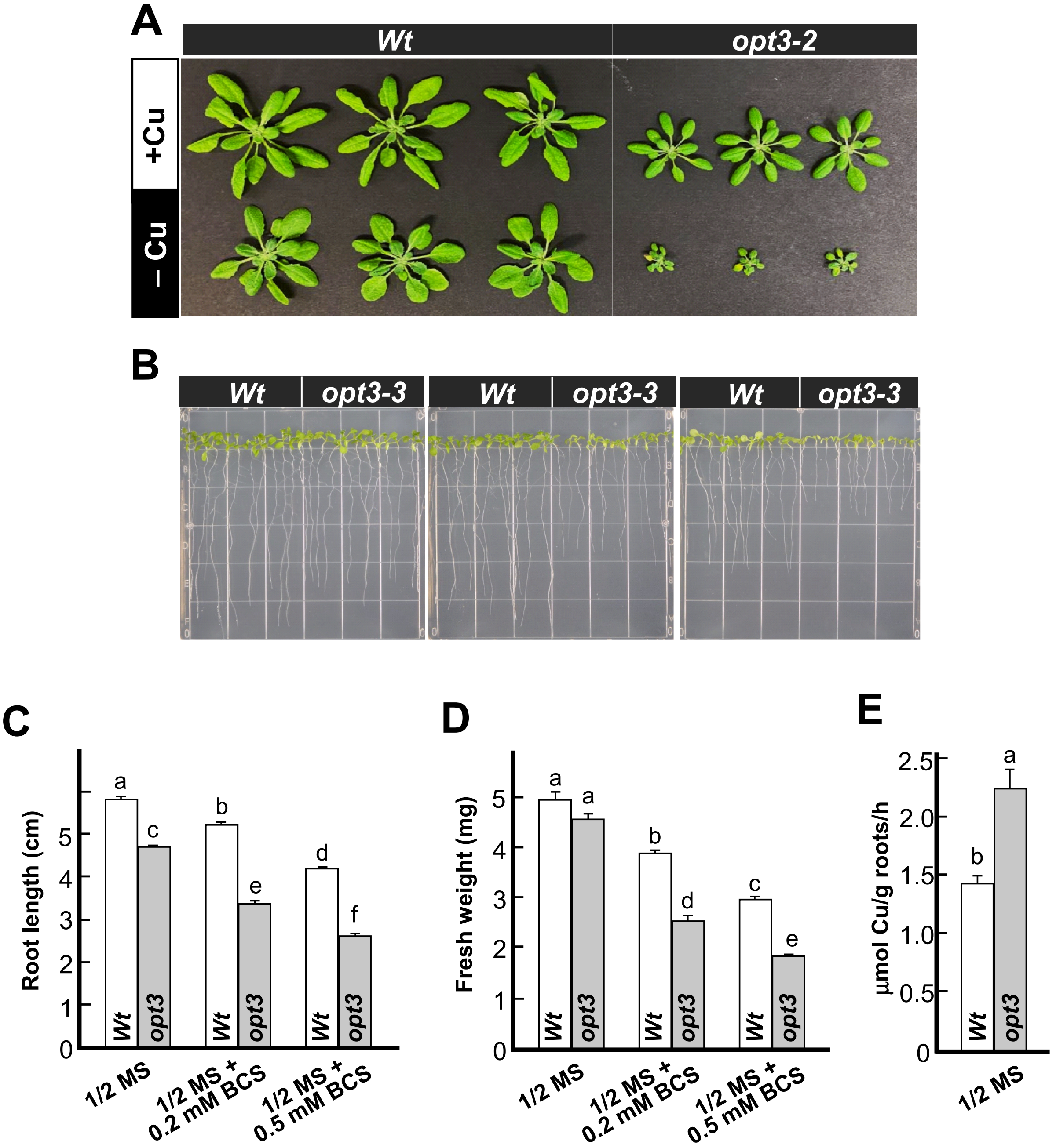


**Supplemental Figure S6 (Supports Fig. 4). Seedlings of the *opt3* mutant are sensitive to copper deficiency.**

(**A**) A representative image of increased sensitivity of the *opt3-2* mutant allele to copper deficiency. Plants were grown hydroponically with or without copper supplements for 35 days. (**B**) to (**E**) The *opt3-3* mutant and wild type were grown on ½ MS solid medium with or without 250 µM or 500 µM of a copper chelator BCS. (**B**) shows a representative image of plants after ten days of growth, after which root length (**C**), fresh weight (**D**) and cupric reductase activity (**E**) were analyzed. In **C** to **E** data are means ± S.E. Levels not connected by the same letter are significantly different ($p < 0.05$, Tukey-Kramer HSD test). In **C**, three independent experiments were performed with 30 plants per genotype analyzed per experiment (n = 90); in **D,** n = 3 independent experiments with 30 to 40 seedlings pooled for analyses in each experiment; in **E**, n = 3 independent experiments with 20 seedlings analyzed in each experiment.

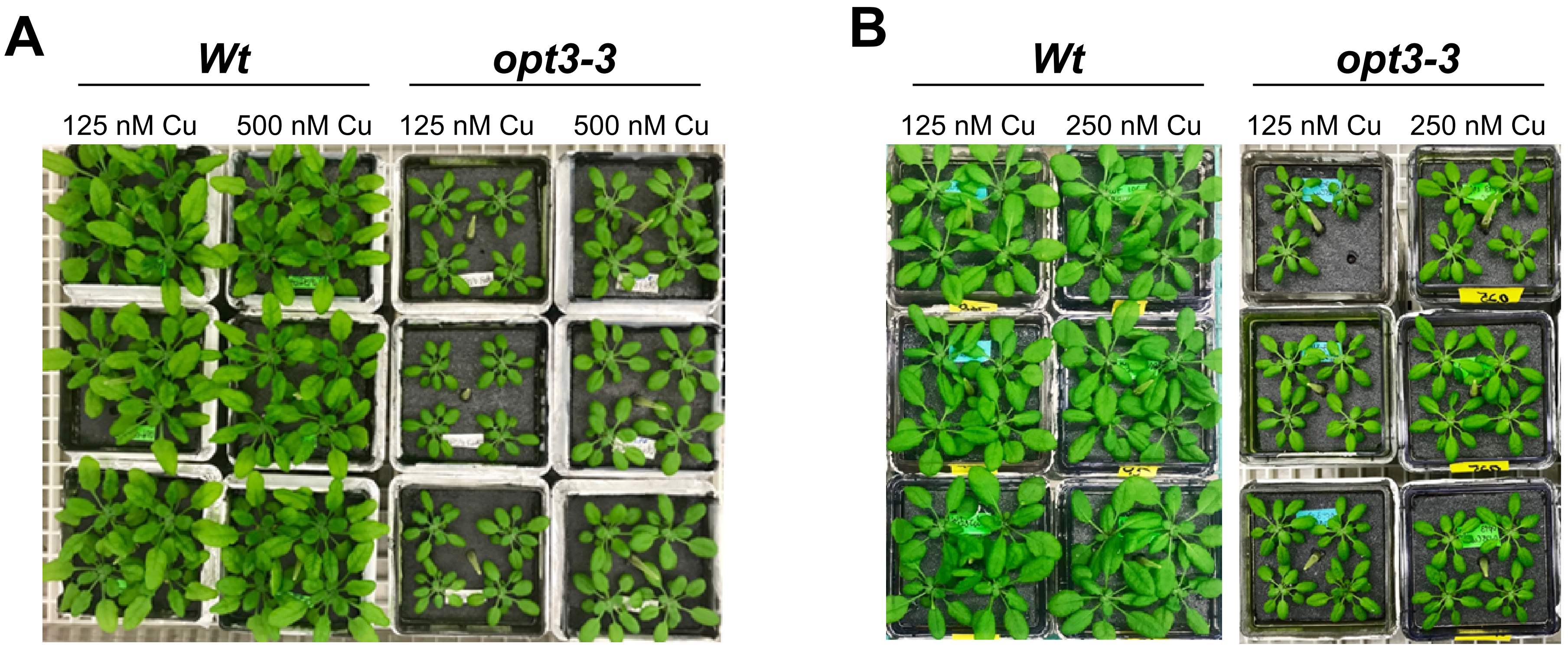


**Supplemental Figure S7 (Supports Fig. 4). Transferring the *opt3-3* mutant to high copper rescues its growth.**

(**A**) Wild type and the *opt3* mutant were germinated and grown hydroponically at 125 nM Cu for four weeks before transferring to a fresh hydroponic medium with 500 nM (**A**) or 250 nM (**B**) copper. Images were taken after one week of growth. A representative set-up/result from three independent experiments is shown.

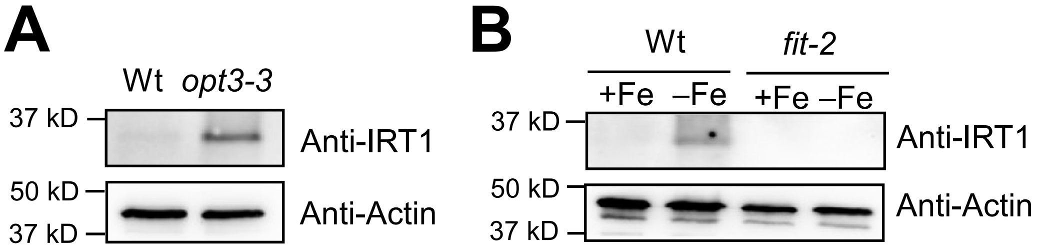


**Supplemental Figure S8 (Supports Fig. 5). The *opt3* mutant accumulates IRT1 protein in roots under Fe sufficient growth conditions.**

(**A**) Western blots analysis of IRT1 protein accumulation in roots of wild type and the *opt3-3* mutant. Plants were grown hydroponically with 125 nM $CuSO_4$ and 10 mM Fe-HBED for four weeks before tissue collection. (**B**) Western blots analysis of iron deficiency-induced IRT1 accumulation in wild type roots. Plants were grown hydroponically with 125 nM $CuSO_4$ and 10 µM Fe-HBED for three weeks and then transferred to a fresh medium without iron. Roots were collected after one week of growth under iron deficiency. The *fit-2* mutant was incorporated as a negative control for the IRT1 protein accumulation. In (**A**) and (**B**), the IRT1 signals are recognized by the antibody at 35 kD while actin (protein loading control) at 45 kD. For the immunodetection of actin epitope, the nitrocellulose blots were probed with the primary mouse-monoclonal anti-actin antibody (1:5,000 dilution, Sigma-Aldrich) and secondary, an HRP-conjugated goat-antimouse IgG antibody (1:10,000 Rockland Immunochemicals). In both cases, immunoreactive bands were visualized with Clarity Max ECL blotting substrates (BIORAD).

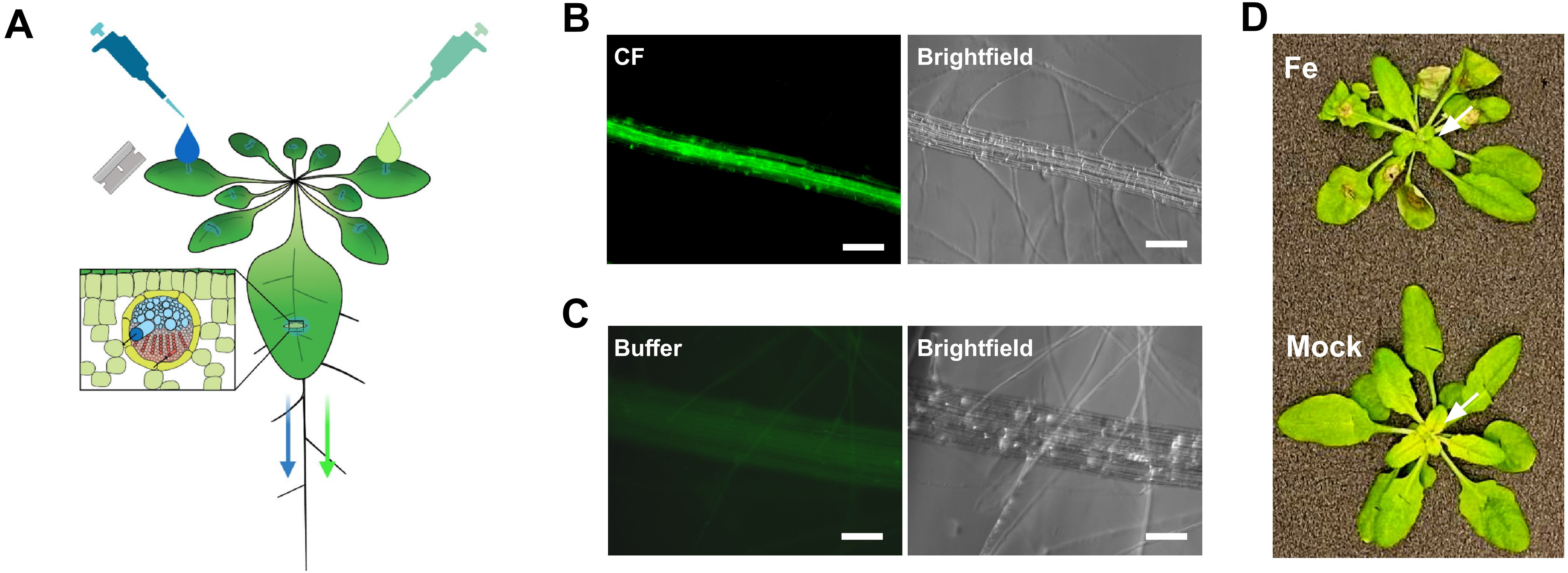


**Supplemental Figure 9 (Supports Fig. 9). Phloem feeding with carboxyfluorescein diacetate succinimidyl ester (CFDA) in leaves of wild type *Arabidopsis thaliana.***

(**A**) The illustration of phloem feeding procedures as described in Materials and Methods. (**B)** Shows a representative image of carboxyfluorescein (**CF**)-mediated fluorescence and brightfield images of the root after feeding 10 mM CFDA *via* the phloem in the shoot. (**C**) Shows autofluorescence of mock-fed plants (**Buffer**). (**D**) shows that iron but not mock feeding via the phloem in the shoot of iron-deficient wild type eliminates degreening of young leaves (indicated by white arrows). In all cases, experiments were repeated at least three times with at least three plasnt analyzed per each experiment.

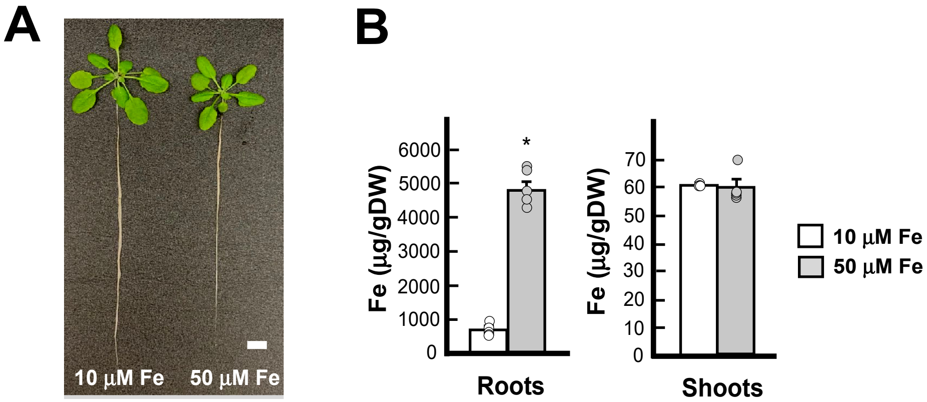


**Supplemental Figure 10 (Supports Fig. 11). Iron availability in the medium influence the expression of copper deficiency responsive genes in the roots of *A. thaliana.***

(**A**) Shows a representative phenotype of *A. thaliana* grown hydroponically with 10 µM or 50 µM FeHBED for five weeks. Twelve plants per condition were analysed in total in three independent experiments. (**B**) The accumulation of iron in roots and shoots of plants grown as in (**A**). Mean values ± S.E (n = 5 ICP-MS measurements obtained from two batches of independently grown plants per condition. Tissues from four plants were pooled for each measurement). Asterisks indicate statistically significant differences ($p < 0.05$, Student-*t* test).
